# Pollen sterols are associated with phylogenetics and environment but not with pollinators

**DOI:** 10.1101/2020.12.18.423259

**Authors:** Pengjuan Zu, Hauke Koch, Orlando Schwery, Samuel Pironon, Charlotte Phillips, Ian Ondo, Iain W. Farrell, W. David Nes, Elynor Moore, Geraldine A. Wright, Dudley I. Farman, Philip C. Stevenson

## Abstract

- Phytosterols are primary plant metabolites that have fundamental structural and regulatory functions. They are also essential nutrients for phytophagous insects, including pollinators, that cannot synthesize sterols. Despite the well-described composition and diversity in vegetative plant tissues, few studies have examined phytosterol diversity in pollen.
- We quantified 25 pollen phytosterols in 122 plant species (105 genera, 51 families) to determine their composition and diversity across plant taxa. We searched literature and databases for plant phylogeny, environmental conditions, and pollinator guilds of the species to examine the relationships with pollen sterols.
- 24-methylenecholesterol, sitosterol and isofucosterol were the most common and abundant pollen sterols. We found phylogenetic clustering of twelve individual sterols, total sterol content and sterol diversity, and of sterol groupings that reflect their underlying biosynthesis pathway (24 carbon alkylation, ring B desaturation). Plants originating in tropical-like climates (higher mean annual temperature, lower temperature seasonality, higher precipitation in wettest quarter) were more likely to record higher pollen sterol content. However, pollen sterol composition and content showed no clear relationship with pollinator guilds.
- Our study is the first to show that pollen sterol diversity is phylogenetically clustered and that pollen sterol content may adapt to environmental conditions.

## Introduction

Phytosterols are a class of lipids with key metabolic and ecological functions for plants (Nes & McKean, 1977; Vanderplanck *et al*., 2020a). For example, they regulate membrane fluidity and permeability (Grunwald, 1971; Schuler *et al*., 1991; Hartmann, 1998), and act as precursors for metabolic signals such as brassinosteroid growth hormones (Grove *et al*., 1979; Chung *et al*., 2010) that promote cell division, and mediate reproduction in plants and protect them against environmental stresses (Khripach *et al*., 2000). Phytosterols may also modulate plant defence against bacterial pathogens (Posé *et al*., 2009; Wang *et al*., 2012; Ferrer *et al*., 2017) and pollen sterols accelerate germination and tube growth and protect against desiccation (Kumar *et al*., 2015; Rotsch *et al*., 2017).

Phytosterols show considerable diversity with more than 250 structures reported (Nes, 2011) with notable variation at the methine substitution (double bond) in ring B and methyl or ethyl substitutions at C-24 (Fig. 1). The structural variation and composition of sterols in plant tissues is important for phytophagous insects since they cannot synthesize sterols *de novo*, and therefore depend upon specific plants to obtain the required sterols from their diet to sustain their development (Behmer & Elias, 1999, 2000; Lang *et al*., 2012). This may be especially important for pollen feeding insects that require specific sterols. Honeybees, for example, require 24-methylenecholesterol (Herbert *et al*. 1980; Chakrabati *et al*., 2020) so must collect pollen from plant species that produce this sterol to rear brood. Bee sterols are similar to those occurring in the pollen on which they feed (Vanderplanck *et al*., 2020a) but differ across bee taxa suggesting bees are what they eat with respect to sterols. Wild pollinators range from pollen generalists to specialists (Rasmussen *et al*., 2020) and for some species, this specialism may be mediated by pollen sterols. Therefore, a landscape of flowers that does not provide the sterols required for a specific bee may be nutritionally deficient for that species. In general, however, the relationships between pollen sterols and the nutritional needs of pollination insects has not yet been evaluated.

**Fig. 1.**
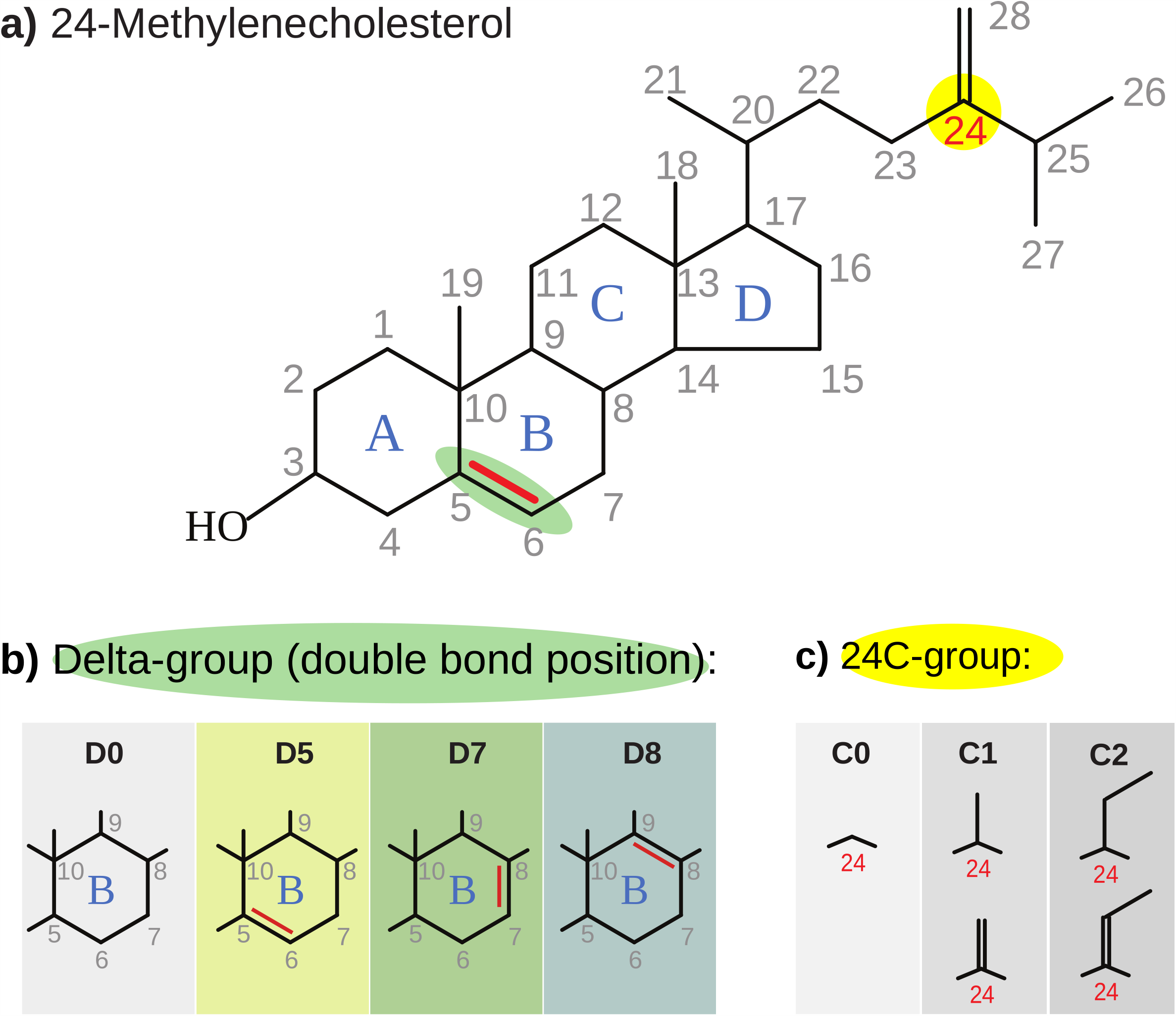
Chemical structure of 24-methylenecholesterol as an illustration of phytosterols showing a) carbon numbering; b) different substitutions in ring B; and c) different substitutions at C-24.

Conversely, plant sterol composition may evolve with antagonists as well as mutualists since the pathways for the synthesis of sterols overlap with that for some defence compounds against herbivores (Qi *et al*., 2006). A range of naturally occurring insect toxins occur in pollen (Arnold *et al*., 2014; Rivest & Forrest, 2019) with the likely role of protecting the male gamete and since some sterols can also act as defensive compounds against arthropod herbivores (Jing & Behmer, 2020). They could also be toxic to pollen feeding insects to reduce damage to or excessive loss of pollen grains.

Abiotic conditions may affect phytosterol structural variations at different levels. At plant individual level, a 24-ethyl substitution (e.g., sitosterol and stigmasterol), for example, reinforces membrane cohesion (Piironen, 2000; Dufourc, 2008), and therefore sterol structures may be altered in response to temperature variations. At population level, from limited heritability studies on phytosterols in plant seeds (Amar *et al*., 2008, Velasco *et al*., 2013), environmental factors also contribute to sterol phenotypic variation, although much less compared to the contribution from genetic factors (heritabilities above 0.8 were documented). At species level, pollen sterol composition seems to be highly variable between different species (Villette *et al*., 2015; Vanderplanck *et al*., 2020a) and can differ from vegetative tissues (Nes, 1990; Nes *et al*., 1993). However, no study has investigated whether pollen sterol variations at species level can be the consequences of evolutionary adaptation to environmental conditions.

Moreover, due to the limited number of studies on pollen sterol profiles, we lack a comprehensive and fundamental understanding of the patterns of pollen sterol diversity across plant taxa. It is still controversial whether pollen phytosterols are phylogenetically structured. For example, Standifer *et al*. (1968) suggested a lack of phylogenetic constraints of pollen sterol composition based on the evidence of large variation in three Salicaceae species. Vanderplanck *et al*. (2020a), in contrast, found similar pollen sterol composition within the genus *Salix* and our interpretation of the data published by Villette *et al*. (2015) suggested the occurrence of some pollen sterols was phylogenetically constrained. Since most studies focused on a few plant species, they were insufficient to reach a general overview of the patterns of pollen sterol diversity across plant taxa and their drivers.

In this study, we analysed pollen sterols including saturated stanols in 122 angiosperms representing 51 plant families and 25 plant orders. We further compiled data from literature and databases on plant phylogeny, pollinators, and environmental conditions within native geographic regions for each plant species to examine relationships between these factors and pollen sterol composition and diversity. Specifically, we ask 1) Are pollen sterols phylogenetically clustered? 2) Are pollen sterols correlated with abiotic environments? 3) Are pollen sterols associated with pollinator guilds?

## Materials and methods

### Pollen collection

From March to November 2018, we collected pollen from fresh flowers growing in the Royal Botanic Gardens (RBG), Kew, UK and nearby areas (see Table S1 for details of collection dates and locations for each species). RBG Kew supports a diverse collection of living plant species from across the world. Prior to pollen collection, we used a fine-meshed bag to cover flower buds whenever possible to prevent potential contamination or removal due to pollinator visitation. When flowers were fully open, we gently shook the flower and collected pollen into a weighed 2 mL microcentrifuge tube (Eppendorf®, Safe-Lock™). For species for which pollen was more difficult to harvest, such as in the cases of *Lamium purpureum* L. and *Ulex europaeus* L., we used small forceps to help push the pollen out or trigger pollen ejection, respectively. Pilot studies carried out in our laboratory (with *Helleborus foetidus, Prunus avium, Prunus spinosa, Salix cinerea* and *Symphytum officinale*) showed a conserved pattern of pollen sterol composition: within species variation was significantly lower than between species variation (all p-values < 0.001 under multi-variate distribution tests e.g., Hotelling test, Pillai test, and Wilks’ lambda distribution test), consistent with findings on within vs. between species variation of other pollen metabolites (Palmer-Young *et al*., 2019). Therefore, we collected 2 to 5 replicates per species (details see Table S1) and used the average quantities across replicates of each species for analyses. In total, we collected 308 samples from 122 species, representing 105 genera, 51 families and 25 orders across the major groups of seed plants (Gymnosperms, Nymphaeales, Monocots, Ranunculids, Caryophyllales, Asterids and Rosids; Table S1). Our selection of species was guided by a combination of practical considerations (feasibility to collect sufficient pollen for analysis, availability of species at Kew) while attempting to maximize phylogenetic and ecological diversity of plants (pollinator guilds, ecological niches). Pollen weight (to 0.1 mg accuracy) and collection date were recorded for each sample. Pollen samples were stored in a freezer (−20°C) before extracting sterols.

### Sterol content analysis

To extract sterols and stanols (from here referred to as phytosterols or pollen sterols) from the pollen, we added 500 μl 10% KOH in MeOH to the microcentrifuge tubes containing a weighed pollen sample. Then, an internal standard (20 μl of 0.5 mg ml^-1^ epicoprostanol) was added prior to incubating the tube for 2 h at 80°C for saponification. Phytosterols were then recovered into 1 mL hexane. After complete evaporation of hexane, phytosterols remained in the tube. We derivatized these with 20 μl Tri-Sil (Sigma, Gillingham, Dorset UK) and then briefly vortexed and injected directly into an Agilent Technologies (Palo Alto, CA, USA) 7890A gas chromatograph connected to an Agilent Technologies 5975C MSD mass spectrometer (GC-MS) and eluted over an Agilent DB5 column using a splitless injection at 250°C with a standard GC program at 170°C for 1 minute ramped to 280°C at 20°C per minute and monitoring between 50 and 550 amu.

All 25 phytosterols were identified by comparison of their retention time relative to cholesterol and mass spectra from authentic standards (David W Nes collection, details see Fig. S4 for mass spectra of each sterol) either directly through co-analysis or using existing data and confirmed where data was available with the NIST (National Institute of Standards and Technology) mass spectral library (Guo *et al*., 1995; Heupel and Nes, 1984; Nes *et al*., 1977; Xu *et al*., 1988; Zhou *et al*., 2009; Nes *et al*., 2003).

To quantify the amount of each phytosterol, we used its relative peak area by calculating the ratio of the peak area of the targeted sterol to that of the internal standard. Then, by multiplying the ratio with the mass of the internal standard, we obtained the mass of each sterol in the sample. Compound identification (using target ion) and quantification were carried out with ChemStation Enhanced Data Analysis (Version E.01.00). In total, we identified 25 phytosterols in pollen (Table S1).

For each plant species, we calculated each phytosterol amount (µg per mg sampled pollen), total sterol content (µg per mg sampled pollen), and the percentage of each sterol in total phytosterol content. In addition, we calculated the chemical diversity index using Shannon entropy: where S is the total number of phytosterols, p^i^ is the percentage of the i^th^ phytosterol. Note that we used the total phytosterol number S as the base of log (instead of the natural base *e*) to scale the range of diversity index values to [0, 1] with 1 indicating the highest diversity. This equates to calculating Shannon’s equitability. Finally, for each phytosterol, we calculated its commonness and abundance across all plant species. Commonness is given by the proportion of plant species that contained that specific phytosterol (i.e., present/absent). Relative abundance was given by the average proportion of a specific sterol across all species.

Additionally, to understand how different phytosterol in pollen co-varied, we performed a factor analysis using the R package *stats* (R Core Team, 2020) on the data for the absolute weight of phytosterols measured in pollen across the entire data set. We set a criterion of eigenvalue > 1 for inclusion of extracted factors. A varimax rotation was used to adjust the fit of the factor analysis to variance in the data.

Moreover, based on biosynthetic reasoning as discussed by Benveniste (2004), we arranged these phytosterols identified in our pollen samples into alternate hypothetical biosynthetic pathways to cholesterol and 24-alkyl phytosterols.

### Phylogenetic tree construction and analyses

We used the R package *rotl* (Michonneau *et al*., 2016) to download the induced subtree of only our focal taxa from the Open Tree of Life (OTL) synthetic tree (Hinchliff *et al*., 2015; Rees *et al*., 2017). If only the genus was known, OTL used the root of the genus for the subtree wherever possible. Name synonyms and corrections suggested by OTL for genus and species were adopted in our analyses (see Table S2). Taxa with subspecies or other epithets beyond species level were reduced to genus and species only (*Amaryllis belladonna* L., *Campanula fragilis* Cirillo, *Campanula isophylla* Moretti, *Euphorbia milii* Des Moul., *Hieracium umbellatum* L.). Only one terminal was retained to represent the two differently coloured varieties of *Hymenocallis littoralis* (Jacq.) Salisb.

We estimated divergence times with penalised likelihood using nine secondary calibration points. Using the R package *ape* (Paradis *et al*., 2004), we randomly resolved polytomies and computed branch lengths using Grafen’s method. We looked up the inferred ages of seven clades from the large phylogeny of spermatophytes by Zanne *et al*., (2014): Angiospermae (243 million years ago [mya]), Monocotyledoneae (171 mya), Eudicotyledoneae (137 mya), Superrosidae (118 mya), Rosidae (117 mya), Superasteridae (117 mya), and Asteridae (108 mya). The age of Nymphaea (78.07 mya) was obtained from DateLife (Sanchez-Reyes, 2019), and we took the estimated origin of Spermatophyta at 327 mya (Smith *et al*., 2010) to calibrate the root age. We used those times as minimal age constraints for a penalized likelihood analysis using *chronopl* in *ape* (Paradis *et al*., 2004). Monophyly of families was checked using *MonoPhy* (Schwery & O’Meara, 2016).

To determine whether there is phylogenetic structure in the pollen sterol data, we used the function *phyloSignal* from the R package *phylosignal* (Keck *et al*., 2016) to calculate Pagel’s λ (Pagel, 1999) and Blomberg’s K (Blomberg *et al*., 2003), each with 999 iterations for *p*-value estimation. Phylogenetic signal was estimated this way for each of the individual sterol compounds (based on their percentage value), for sums of compounds belonging to each C-24 substitution (C0, C1, C2 indicating substitution with no carbon, a methyl and an ethyl), and position of the olefinic or methine moiety in ring B (Δ^0^, Δ^5^, Δ^7^, Δ^8^), for the sterol diversity index H, and for the total phytosterol content (absolute sterol amount per mg pollen). The output of these analyses was visualized using the R packages *phytools* (Revell 2012) and *picante* (Kembel *et al*. 2010).

### Plant occurrence records and abiotic environmental data

To investigate whether species-level variations in pollen sterols are partially the consequences of evolutionary adaptation to environmental conditions, we retrieved environmental information of the native geographic ranges of each species. Note that here we focused on “long-term” prevailing abiotic conditions (e.g., climate) capable of shaping evolutionary changes of sterol composition at species level, as opposed to “short-term” abiotic variables (e.g., stresses, weather) affecting traits via phenotypic plasticity at the individual level. For each species, we extracted geographic occurrence records from several global and continental databases: GBIF (Global Biodiversity Information Facility; https://www.gbif.org/) using the *rgbif* package in R, BIEN (Botanical Information and Ecology Network; http://bien.nceas.ucsb.edu/bien/) using the *BIEN* R package, BioTIME (Dornelas *et al*., 2018) and Rainbio (Dauby *et al*., 2016). Because raw occurrence data from these databases contain taxonomic, spatial and temporal inconsistencies (Meyer *et al*., 2016), we applied different cleaning filters using the *CoordinateCleaner* package in R (Zizka *et al*., 2019). We discarded non-georeferenced records, records with latitude and longitude given as zero and having equal longitude and latitude, points recorded before 1950, as well as fossil data, records corresponding to centroids of countries, capitals, known botanical institutions and GBIF headquarters, occurrences falling in the sea, cultivated records, and points indicated as having high coordinate uncertainty (>20 km). We used the World Geographical Scheme for Recording Plant Distribution (WGSRPD) database (Brummitt, 2001) from the World Checklist of Vascular Plants (WCVP, 2020) to discard records from species reported outside of their native regions at the level-2 (regional or sub-continental level). Finally, we removed duplicates and thinned each species’ occurrence dataset by keeping only one record by 10×10 arc-min grid cell to limit spatial autocorrelation. In total, 355,912 occurrence records were retrieved across all species (Table S1).

We quantified species environmental niches based on a set of 13 climate, soil, and topography variables. Eight of them were bioclimatic variables (BIO1, BIO4, BIO10, BIO11, BIO12, BIO15, BIO16 and BIO17) extracted from the CHELSA database (Karger *et al*., 2017), representing annual mean, seasonality, minimum and maximum temperature and precipitation (full list of variables and descriptions see Table S3). Four soil variables were extracted from the SoilGrids database (ISRIC, 2013; http://www.data.isric.org) and averaged across a 0-60 cm depth gradient: depth to bedrock, mean soil organic carbon stock, pH and water capacity. Land slope was calculated using the Slope function in the Spatial Analyst toolbox of ARC/INFO GIS based on the Global Multi-resolution Terrain Elevation Database (GMTED) (Danielson *et al*., 2011). To match the resolution of the occurrence records, all environmental variables were upscaled to 10 arc-min (ca. 20 km) using the *resample* function of the *raster* package in R.

We extracted each of the 13 environmental variables at each occurrence point of each species using the *extract* function of the *raster* package in R. Mean environmental conditions were then calculated for each of the 13 variables across all occurrences of each species (i.e., environmental niche position along individual environmental gradients). We also created an environmental space summarizing the variation in the 13 environmental variables across the world using a Principal Component Analysis (PCA) and the function *princomp* in the *stats* package in R. We kept the first three component axes that explained 74% of the variation in the 13 variables: PC1 being mainly positively correlated with mean temperatures and negatively correlated with temperature seasonality and soil carbon content, PC2 being positively correlated with soil pH and negatively with precipitation, and PC3 being positively correlated with soil depth to bedrock and negatively with land slope (see Table S3 for variable contributions to PCA axes). To quantify the niche breadth of each species, we first drew three-dimensional alpha shapes around each set of occurrence points of each species in the environmental space defined by the PCA with an alpha value of two using the *ashape3d* function in the *alphashape3d* package in R (Capinha & Pateiro-López, 2014). The alpha-shape is a profile technique used to compute species environmental niche envelopes using a flexible envelope fitting procedure that does not make any assumption about the shape of the niche (Capinha & Pateiro-López, 2014). We then calculated the volume of each species’ alpha shape as a measure of their environmental niche breadth using the *volume_ashape3d* function from the latter package. We also calculated the mean position of each species’ alpha shape on the three retained main axes of the PCA (i.e., niche position, individual variable contributions see Table S3). Because three-dimensional alpha shapes require at least five occurrence points to be drawn, species with fewer records were discarded. We also discarded those species lacking sufficient and reliable geographic data or taxonomic uncertainty (i.e., we did not extract occurrence records for genera, subspecies and hybrids). In the end, we quantified niche breadth for 90 species, while 32 taxa were discarded, and niche position for 100 species (22 taxa discarded; details see Table S1).

### Pollinator data collection

To study whether there is a relationship between plants’ pollen sterols and their pollinators, we categorized plants in two different ways. Firstly, based on pollinator guilds, as 1) Bee, 2) Fly, 3) Lepidoptera, 4) Thrips, 5) Generalist insect, 6) Bird, or 7) Wind pollinated. Secondly, we grouped plants by whether or not pollen acts as a reward for bee pollinators. On the one hand, bees depend on pollen as larval food and require pollen sterols as essential nutrients. Plants could therefore hypothetically attract bee pollinators with pollen sterol profiles of high nutritional quality to them. On the other hand, if pollen does not play a role as bee reward (i.e., in non-bee pollinated plants, and/or where nectar is the sole reward), sterol profiles could be expected that are of low quality or even toxic to bees to prevent pollen robbery (as shown for some other chemical compounds in pollen, Rivest & Forrest 2019).

To classify plant pollinator guilds and groups, we conducted literature searches for each plant species on Google Scholar, using the scientific name (including common synonyms) and “pollinat*”, OR “pollen”, OR “flower” as search terms. We examined relevant cited or citing references of publications found in this way for additional records, and consulted Knuth (1908, 1909) and Westrich (2018), or personal observations. If no sources on pollination and flower visitation were available, the pollinator guild was classified as “unknown” (10 species in data set). We included plant species in the “pollen as bee reward” group that both receive pollination services by bees (including some plants in the “generalist insect pollination” category) and have records of bees collecting pollen. Plants were classified as not producing pollen as bee reward if they were either not pollinated by bees, or, in case of bee pollination, had clear evidence of pollen not being collected by bees (e.g., pollen contained in pollinia of bee-pollinated orchids). Plants for which data on pollinator guild and collection of pollen by bees was missing were classified as “unknown” (34 species in data set). A full list of relevant references and the assigned pollinator guilds is provided in Table S1.

### Analyses on relationships between phytosterols and (a)biotic factors

To assess the association of sterol composition with environmental variables and pollinator guilds, we first calculated a Bray-Curtis distance matrix for sterol profiles of pairwise plant species comparisons, using absolute weights (µg) of each sterol per mg pollen. Then we related this distance matrix to environmental factors and pollinator guild to study their relationships. Specifically, for abiotic environmental factors (continuous values), we ran MRM (multiple regression on distance matrices) analyses (Lichstein, 2007) using an additive linear model with the Bray-Curtis distance matrix of pollen sterol composition dissimilarity as response, and environmental niche distance matrices for PC1, PC2, PC3 (see “abiotic environmental data” above, calculated from pairwise Euclidean distances for all plants for their position on each of the PCs) and a phylogenetic distance matrix (pairwise phylogenetic distance in mya, phylogeny see above) as independent variables. We used Pearson correlations with 10000 permutations to test for significant associations. Calculations of distance matrices and MRM analyses were done with the R package *ecodist* (Goslee & Urban, 2007). For pollinator modes (categorical variables), we conducted ANOSIMs (analyses of similarities; Clarke, 1993) to test for significant differences of pollen sterol profiles between different pollinator groups (excluding pollinator groups with only one representative, i.e. wind, thrips, and fly) or between plant groups where pollen is used as reward by bees or not. ANOSIMs were conducted in PAST 4.03 (Hammer *et al*., 2001) with 10000 permutations. We illustrated the relationship of these factors to sterol profile similarity with 2D non-metric multidimensional scaling (NMDS) ordination plots in PAST 4.03 based on Bray-Curtis dissimilarities.

We furthermore examined associations of environmental variables and niche breadth with total sterol content and diversity. We calculated phylogenetic independent contrasts (Felsenstein, 1985; implemented in R package *ape* (Paradis *et al*., 2004)) with the phylogeny outlined above for sterol contents, Shannon diversity H, positions on environmental principal component axes (PC1, PC2, PC3), and environmental niche breadth. Associations between contrasts of sterol content or diversity (as dependent variable) with contrasts of environmental principal components, niche breadth, or the 13 separate environmental factors were then individually evaluated by linear models in R (fitting the regression through the origin).

Finally, as 24-methylenecholesterol is a key sterol nutrient for honey bee development (Svoboda *et al*., 1980; Herbert *et al*., 1980), and could therefore have been selected for as an attracting reward in bee pollinated plants, we tested for differences in 24-methylenecholesterol content for plants that offer pollen as reward for bees or not (or for which this interaction was unknown) with a phylogenetic ANOVA (Garland *et al*., 1993), implemented in the R package *phytools* (Revell, 2012), with 1000 simulations, and “Holm” post-hoc testing. The same test was also conducted for total sterol content. Only species with phylogenetic information were included (pollen as bee reward: n = 54; pollen not bee reward: n = 22; unknown: n = 24).

## Results

### Pollen sterol composition and diversity across taxa

We profiled 25 phytosterols in pollen of 122 plant species from 51 families including representatives of Gymnosperms, Nymphaeales, Monocots, Ranunculids, Caryophyllales, Asterids and Rosids (Fig. 2, Table S1). These phytosterols can be arranged into biosynthetic pathways with three main distinct branches (i.e., 24C-0, 24C-methyl and 24C-ethyl groups, Fig 3. See Fig. 1 for structure-illustration of the groups). Pollen phytosterols varied qualitatively and quantitatively across taxa with each species exhibiting a distinctive sterol profile (Fig. 2).

**Fig. 2.**
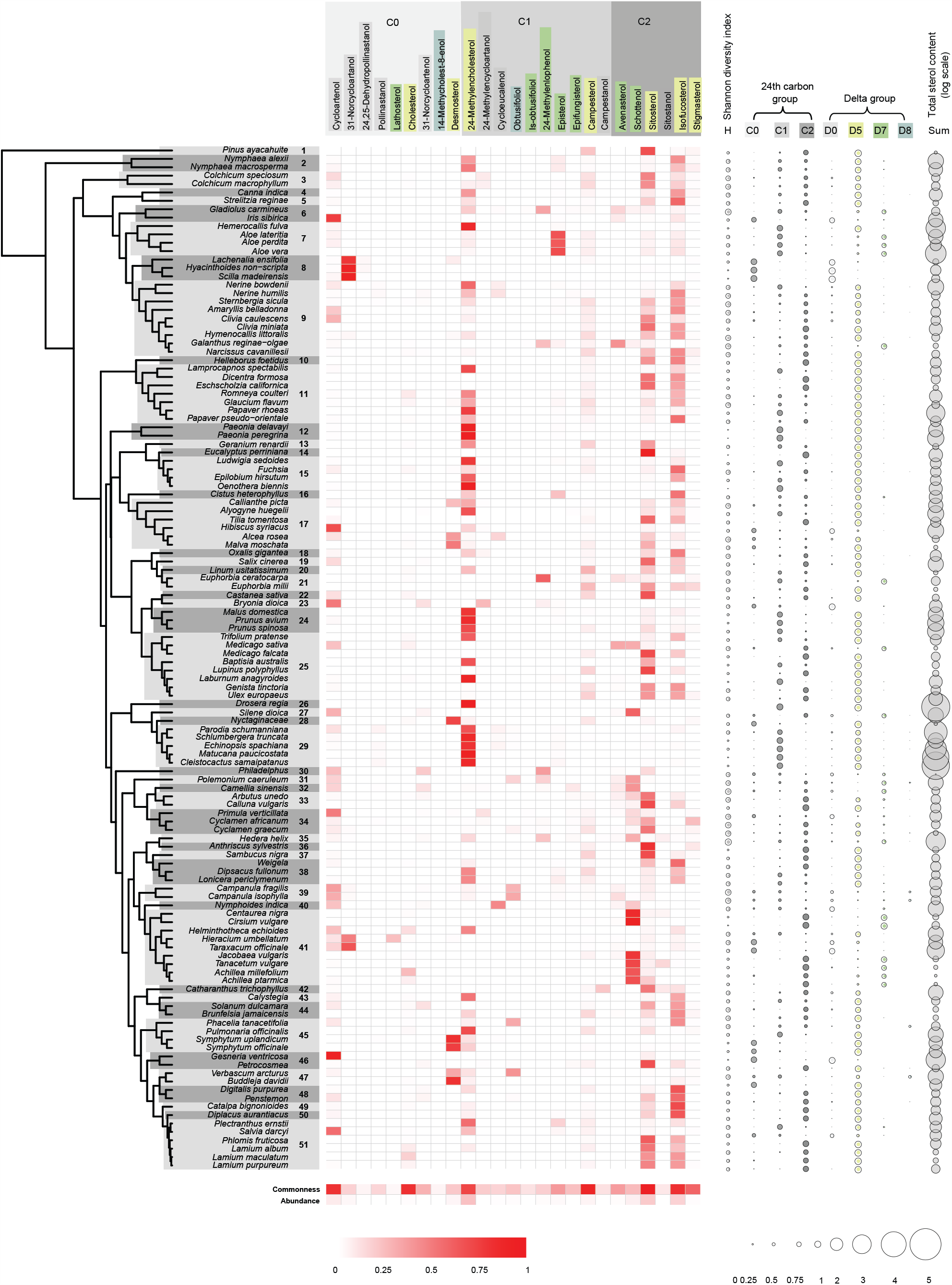
Pollen sterol profiles of plant species. Phylogenetic relationships are given on the left, bold numbers indicate families. Relative contributions of individual sterols to each species’ total sterol content are given in the centre; commonness (proportion of plants containing an individual sterol) and relative abundance (average proportion of individual sterol in each species) are given at the bottom; deeper reds indicate values closer to 1. Shannon diversity index (H), 24th carbon groups, delta groups, and total sterol content are given on the right; circle size represents sums of relative sterol contents in the respective groups (0 to 1), and log of µg per mg pollen for total sterol content. Sterol names and groups are coloured in the same fashion as illustrated in Fig. 1. Families: 1 - Pinaceae, 2 - Nymphaeaceae, 3 - Colchicaceae, 4 - Cannaceae, 5 - Strelitziaceae, 6 - Iridaceae, 7 - Asphodelaceae, 8 - Asparagaceae, 9 - Amaryllidaceae, 10 - Ranunculaceae, 11 - Papaveraceae, 12 - Paeoniaceae, 13 - Geraniaceae, 14 - Myrtaceae, 15 - Onagraceae, 16 - Cistaceae, 17 - Malvaceae, 18 - Oxalidaceae, 19 - Salicaceae, 20 - Linaceae, 21 - Euphorbiaceae, 22 - Fagaceae, 23 - Cucurbitaceae, 24 - Rosaceae, 25 - Fabaceae, 26 - Droseraceae, 27 - Caryophyllaceae, 28 - Nyctaginaceae, 29 - Cactaceae, 30 - Hydrangeaceae, 31 - Polemoniaceae, 32 - Theaceae, 33 - Ericaceae, 34 - Primulaceae, 35 - Araliaceae, 36 - Apiaceae, 37 - Adoxaceae, 38 - Caprifoliaceae, 39 - Campanulaceae, 40 - Menyanthaceae, 41 - Asteraceae, 42 - Apocynaceae, 43 - Convolvulaceae, 44 - Solanaceae, 45 - Boraginaceae, 46 - Gesneriaceae, 47 - Scrophulariaceae, 48 - Plantaginaceae, 49 - Bignoniaceae, 50 - Phrymaceae, 51 - Lamiaceae.

**Fig. 3.**
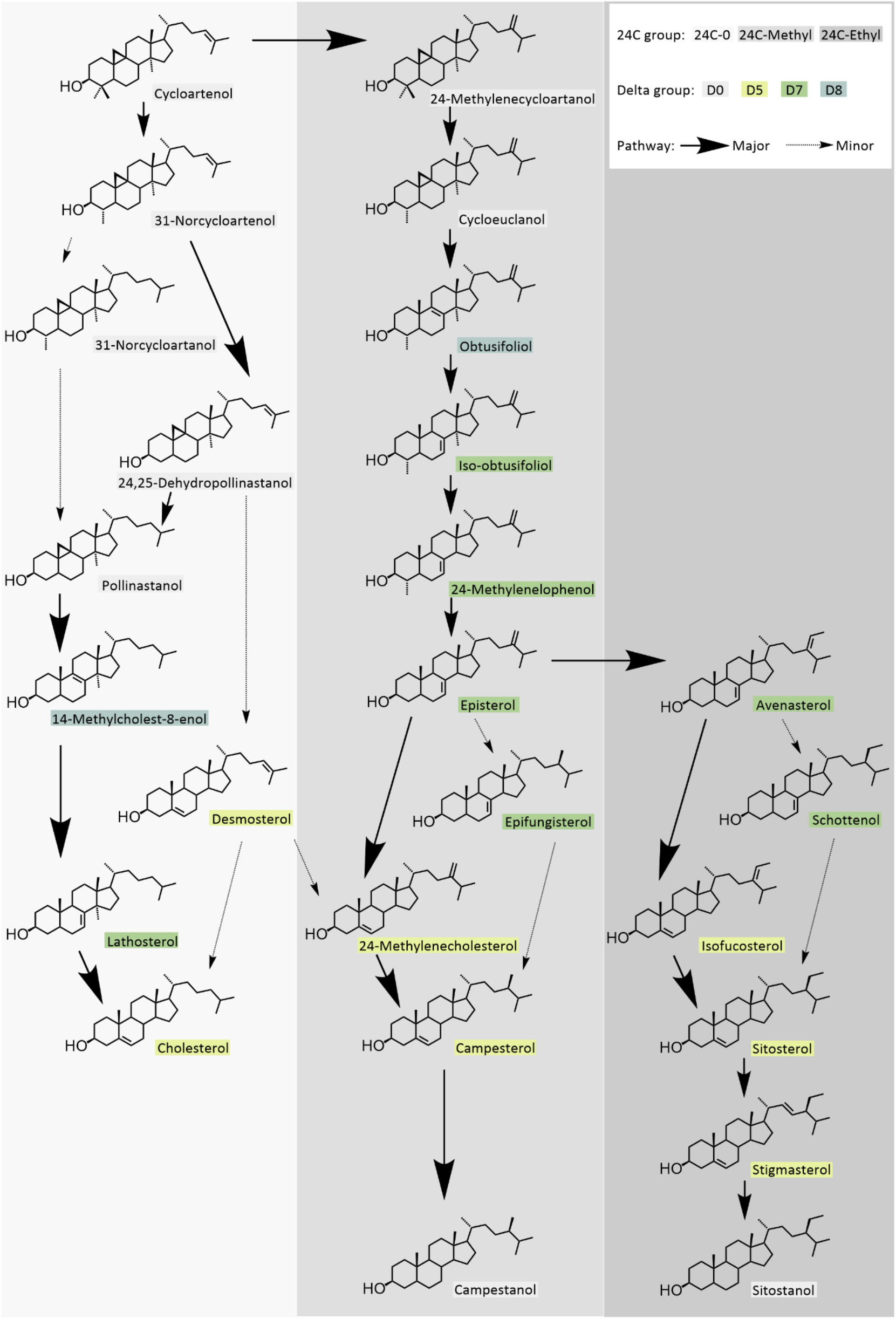
Hypothetical biosynthetic pathways of phytosterols identified in this study (pathways based on Benveniste, 2004).

Across all the sampled species, the most common pollen sterols (labelled “commonness”, Fig. 2) were sitosterol (recorded in 97.5% sampled species), campesterol (88.5%), isofucosterol (82.0%), cholesterol (82.0%), cycloartenol (81.1%), 24-methylenecholesterol (ostreasterol) (73.0%) and stigmasterol (59.0%). The most abundant sterol dominating pollen sterol profiles (labelled “abundance”, Fig. 2) was 24-methylenecholesterol (on average accounting for 23% of total sterol content), followed by isofucosterol (21.5%), sitosterol (20.7%), and cycloartenol (17.7%). The first three are all Δ^5^ sterols, of which 24-methylenecholesterol belongs to the 24C-methyl group, whereas sitosterol and isofucosterol belong to the 24C-ethyl group. Cholesterol, the primary sterol in animals, only represented a small portion (<1%) of pollen sterol content, despite being common.

The pollen sterol diversity of plants varied dramatically with a mean of 9.98 ± 4.46 (mean ± s.d.) different phytosterols. For example, the carnivorous plant *Drosera regia* Stephens had almost exclusively 24-methylenecholesterol in pollen, whereas pollen from ivy (*Hedera helix* L.) contained 23 different sterols, tea pollen (*Camellia sinensis* L.) had 22 sterols, and pollen from the bellflowers *Campanula fragilis* Cirillo and *Campanula isophylla* Moretti had 23 and 19 sterols respectively. However, in all these species, only one to two sterol compounds were typically major components (contributing >50% of total sterol content). The variation in the total weights of sterols led to a Shannon diversity index for pollen sterol composition ranging from 0 in *Drosera regia* to 0.64 in *Hedera helix* (Fig. 2, Table S1), with a mean of 0.34 (note that we standardized the maximum value of the Shannon diversity index to be 1.0, details see method).

### Covariance of pollen sterols

The factor analysis reduced the data to 12 independent latent factors that explained 73% of sterol variation (Table 1). Overall, phytosterols that have close positions in their biosynthetic pathways (Fig. 3) or use the same enzyme (e.g., reductase) for production tend to align together with the same factors. For example, iso-obtusifoliol is the precursor of 24-methylenelophenol, then it branches to either epifungisterol or avenasterol via episterol (Fig. 3). These four sterol compounds (not including episterol) largely aligned together with factor 1 which accounted for ∼9% of the variance (Table 1). Similar patterns also applied to factor 3 and factor 4 whose main contributing sterols represented the early cyclopropyl pathway intermediates. Factor 5 represented a strong positive correlation among the stanols (saturated in ring B), campestanol and sitostanol. Factors 6 and 7 represent products of Δ-24 reduction. In addition, we found one inverse relationship between four of the most common phytosterols (in factor 2, accounting for 8% of the variance), where 24-methylenecholesterol was aligned in the opposite direction as the presence of three other phytosterols: sitosterol, campesterol and stigmasterol.

**Table 1.**
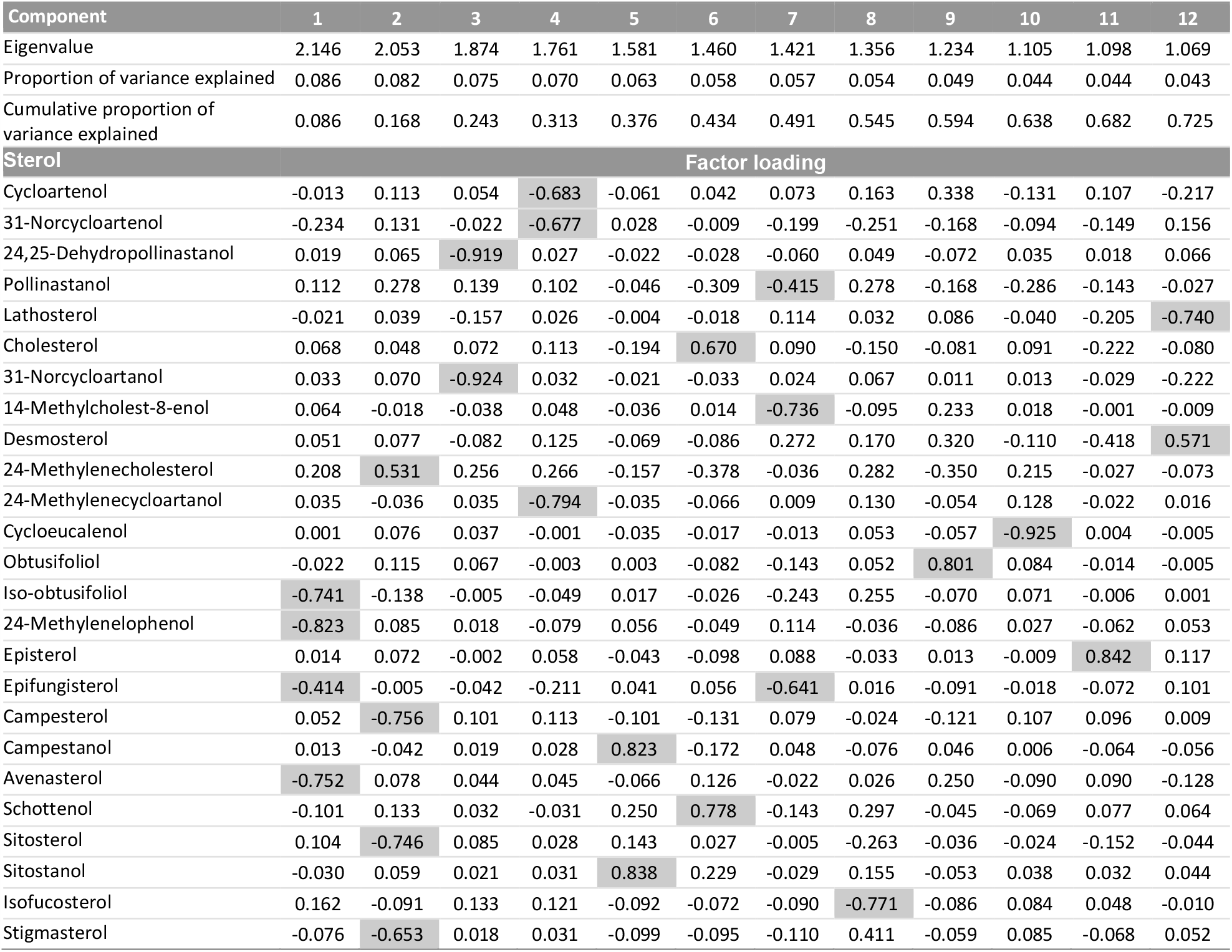
Factor analysis identifying the covariance of 25 sterols measured across all the plant species surveyed. The main contributor(s) for each component is highlighted in grey.

| Component | 1 | 2 | 3 | 4 | 5 | 6 | 7 | 8 | 9 | 10 | 11 | 12 |
| --- | --- | --- | --- | --- | --- | --- | --- | --- | --- | --- | --- | --- |
| Eigenvalue | 2.146 | 2.053 | 1.874 | 1.761 | 1.581 | 1.460 | 1.421 | 1.356 | 1.234 | 1.105 | 1.098 | 1.069 |
| Proportion of variance explained | 0.086 | 0.082 | 0.075 | 0.070 | 0.063 | 0.058 | 0.057 | 0.054 | 0.049 | 0.044 | 0.044 | 0.043 |
| Cumulative proportion of variance explained | 0.086 | 0.168 | 0.243 | 0.313 | 0.376 | 0.434 | 0.491 | 0.545 | 0.594 | 0.638 | 0.682 | 0.725 |
| **Sterol** | **Factor loading** |  |  |  |  |  |  |  |  |  |  |  |
| Cycloartenol | -0.013 | 0.113 | 0.054 | -0.683 | -0.061 | 0.042 | 0.073 | 0.163 | 0.338 | -0.131 | 0.107 | -0.217 |
| 31-Norcycloartenol | -0.234 | 0.131 | -0.022 | -0.677 | 0.028 | -0.009 | -0.199 | -0.251 | -0.168 | -0.094 | -0.149 | 0.156 |
| 24,25-Dehydropollinastanol | 0.019 | 0.065 | -0.919 | 0.027 | -0.022 | -0.028 | -0.060 | 0.049 | -0.072 | 0.035 | 0.018 | 0.066 |
| Pollinastanol | 0.112 | 0.278 | 0.139 | 0.102 | -0.046 | -0.309 | -0.415 | 0.278 | -0.168 | -0.286 | -0.143 | -0.027 |
| Lathosterol | -0.021 | 0.039 | -0.157 | 0.026 | -0.004 | -0.018 | 0.114 | 0.032 | 0.086 | -0.040 | -0.205 | -0.740 |
| Cholesterol | 0.068 | 0.048 | 0.072 | 0.113 | -0.194 | 0.670 | 0.090 | -0.150 | -0.081 | 0.091 | -0.222 | -0.080 |
| 31-Norcycloartanol | 0.033 | 0.070 | -0.924 | 0.032 | -0.021 | -0.033 | 0.024 | 0.067 | 0.011 | 0.013 | -0.029 | -0.222 |
| 14-Methylcholest-8-enol | 0.064 | -0.018 | -0.038 | 0.048 | -0.036 | 0.014 | -0.736 | -0.095 | 0.233 | 0.018 | -0.001 | -0.009 |
| Desmosterol | 0.051 | 0.077 | -0.082 | 0.125 | -0.069 | -0.086 | 0.272 | 0.170 | 0.320 | -0.110 | -0.418 | 0.571 |
| 24-Methylencholesterol | 0.208 | 0.531 | 0.256 | 0.266 | -0.157 | -0.378 | -0.036 | 0.282 | -0.350 | 0.215 | -0.027 | -0.073 |
| 24-Methylenecycloartanol | 0.035 | -0.036 | 0.035 | -0.794 | -0.035 | -0.066 | 0.009 | 0.130 | -0.054 | 0.128 | -0.022 | 0.016 |
| Cycloeucalenol | 0.001 | 0.076 | 0.037 | -0.001 | -0.035 | -0.017 | -0.013 | 0.053 | -0.057 | -0.925 | 0.004 | -0.005 |
| Obtusifoliol | -0.022 | 0.115 | 0.067 | -0.003 | 0.003 | -0.082 | -0.143 | 0.052 | 0.801 | 0.084 | -0.014 | -0.005 |
| Iso-obtusifoliol | -0.741 | -0.138 | -0.005 | -0.049 | 0.017 | -0.026 | -0.243 | 0.255 | -0.070 | 0.071 | -0.006 | 0.001 |
| 24-Methylenelophenol | -0.823 | 0.085 | 0.018 | -0.079 | 0.056 | -0.049 | 0.114 | -0.036 | -0.086 | 0.027 | -0.062 | 0.053 |
| Episterol | 0.014 | 0.072 | -0.002 | 0.058 | -0.043 | -0.098 | 0.088 | -0.033 | 0.013 | -0.009 | 0.842 | 0.117 |
| Epifungisterol | -0.414 | -0.005 | -0.042 | -0.211 | 0.041 | 0.056 | -0.641 | 0.016 | -0.091 | -0.018 | -0.072 | 0.101 |
| Campesterol | 0.052 | -0.756 | 0.101 | 0.113 | -0.101 | -0.131 | 0.079 | -0.024 | -0.121 | 0.107 | 0.096 | 0.009 |
| Campestanol | 0.013 | -0.042 | 0.019 | 0.028 | 0.823 | -0.172 | 0.048 | -0.076 | 0.046 | 0.006 | -0.064 | -0.056 |
| Avenasterol | -0.752 | 0.078 | 0.044 | 0.045 | -0.066 | 0.126 | -0.022 | 0.026 | 0.250 | -0.090 | 0.090 | -0.128 |
| Schottenol | -0.101 | 0.133 | 0.032 | -0.031 | 0.250 | 0.778 | -0.143 | 0.297 | -0.045 | -0.069 | 0.077 | 0.064 |
| Sitosterol | 0.104 | -0.746 | 0.085 | 0.028 | 0.143 | 0.027 | -0.005 | -0.263 | -0.036 | -0.024 | -0.152 | -0.044 |
| Sitostanol | -0.030 | 0.059 | 0.021 | 0.031 | 0.838 | 0.229 | -0.029 | 0.155 | -0.053 | 0.038 | 0.032 | 0.044 |
| Isofucosterol | 0.162 | -0.091 | 0.133 | 0.121 | -0.092 | -0.072 | -0.090 | -0.771 | -0.086 | 0.084 | 0.048 | -0.010 |
| Stigmasterol | -0.076 | -0.653 | 0.018 | 0.031 | -0.099 | -0.095 | -0.110 | 0.411 | -0.059 | 0.085 | -0.068 | 0.052 |

### Phylogenetic patterns

We found significant phylogenetic signal in 12 out of 25 phytosterols (percentages of individual compounds), of which 7 were significant for both Pagel’s λ and Blomberg’s K, and 5 for only one of the tests (Fig.2, Table 2). When grouping phytosterols based on the substitution at C-24 (24C-methyl-, 24C-ethyl-, or 24C-0) or based on the position of methine in ring B (Δ^0^, Δ^5^, Δ^7^, Δ^8^), we found a significant phylogenetic signal (both Pagel’s λ and Blomberg’s K) for all groups except the Δ^8^ sterols (Fig.2, Table 2). Additionally, we found a significant signal for the Shannon diversity index and total sterol content (µg sterol per mg pollen; Fig.2, Table 2). These results remain largely consistent when excluding all taxa which are only identified to genus level. Note that λ and K are largely agreeing on which phytosterols showed significant signal (Table 2), although the significant estimates for λ are relatively high (0.585 to 1, mean = 0.847 for individual compounds; 0.668 to 0.906, mean = 0.79 for categories), whereas those for K are comparatively low (0.183 to 0.505, mean = 0.332 for individual compounds; 0.158 to 0.201, mean = 0.182 for categories). Some phylogenetic clustering of plants by overall sterol compositional similarity was also apparent in the NMDS plot, with for example plants in the Asteraceae, Asparagaceae, or Cactaceae sharing similar sterol profiles (Fig. S1).

**Table 2.**
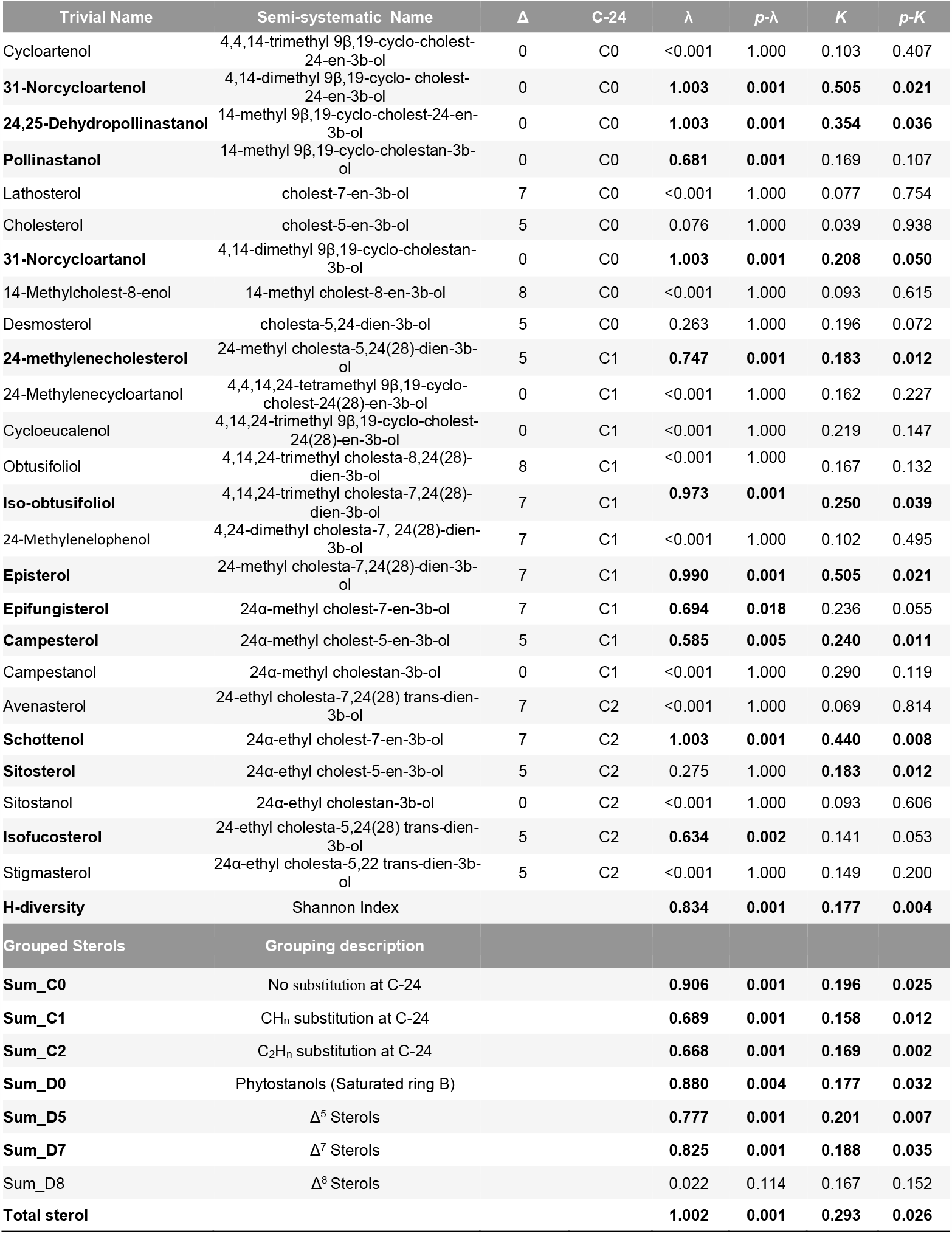
Identity of phytosterols in pollen of 122 plant species showing those with phylogenetic signal across species. All values presented are based on the percentage values of sterols except total sterols content (µg/mg sampled pollen). Δ and C-24 value indicates structure of ring B and on the 24t^h^ carbon (see Fig. 1 for details). Pagel’s λ and Blomberg’s K are used for testing phylogenetic signal. P-values for each test are given accordingly (p-λ and p-K). Sterols with significant phylogenetic signals are in bold.

| Trivial Name | Semi-systematic Name | $\Delta$ | C-24 | $\lambda$ | $p\text{-}\lambda$ | $K$ | $p\text{-}K$ |
| --- | --- | --- | --- | --- | --- | --- | --- |
| Cycloartenol | 4,4,14-trimethyl 9β,19-cyclo-cholest-24-en-3b-ol | 0 | C0 | <0.001 | 1.000 | 0.103 | 0.407 |
| **31-Norcycloartenol** | 4,14-dimethyl 9β,19-cyclo-cholest-24-en-3b-ol | 0 | C0 | **1.003** | **0.001** | **0.505** | **0.021** |
| **24,25-Dehydropollinastanol** | 14-methyl 9β,19-cyclo-cholest-24-en-3b-ol | 0 | C0 | **1.003** | **0.001** | **0.354** | **0.036** |
| **Pollinastanol** | 14-methyl 9β,19-cyclo-cholestan-3b-ol | 0 | C0 | **0.681** | **0.001** | 0.169 | 0.107 |
| Lathosterol | cholest-7-en-3b-ol | 7 | C0 | <0.001 | 1.000 | 0.077 | 0.754 |
| Cholesterol | cholest-5-en-3b-ol | 5 | C0 | 0.076 | 1.000 | 0.039 | 0.938 |
| **31-Norcycloartanol** | 4,14-dimethyl 9β,19-cyclo-cholestan-3b-ol | 0 | C0 | **1.003** | **0.001** | **0.208** | **0.050** |
| 14-Methylcholest-8-enol | 14-methyl cholest-8-en-3b-ol | 8 | C0 | <0.001 | 1.000 | 0.093 | 0.615 |
| Desmosterol | cholesta-5,24-dien-3b-ol | 5 | C0 | 0.263 | 1.000 | 0.196 | 0.072 |
| **24-methylencholesterol** | 24-methyl cholesta-5,24(28)-dien-3b-ol | 5 | C1 | **0.747** | **0.001** | **0.183** | **0.012** |
| 24-Methylenecycloartanol | 4,4,14,24-tetramethyl 9β,19-cyclo-cholest-24(28)-en-3b-ol | 0 | C1 | <0.001 | 1.000 | 0.162 | 0.227 |
| Cycloeucalenol | 4,14,24-trimethyl 9β,19-cyclo-cholest-24(28)-en-3b-ol | 0 | C1 | <0.001 | 1.000 | 0.219 | 0.147 |
| Obtusifolol | 4,14,24-trimethyl cholesta-8,24(28)-dien-3b-ol | 8 | C1 | <0.001 | 1.000 | 0.167 | 0.132 |
| **Iso-obtusifolol** | 4,14,24-trimethyl cholesta-7,24(28)-dien-3b-ol | 7 | C1 | **0.973** | **0.001** | **0.250** | **0.039** |
| 24-Methylenelophenol | 4,24-dimethyl cholesta-7, 24(28)-dien-3b-ol | 7 | C1 | <0.001 | 1.000 | 0.102 | 0.495 |
| **Episterol** | 24-methyl cholesta-7,24(28)-dien-3b-ol | 7 | C1 | **0.990** | **0.001** | **0.505** | **0.021** |
| **Epifungisterol** | 24α-methyl cholest-7-en-3b-ol | 7 | C1 | **0.694** | **0.018** | 0.236 | 0.055 |
| **Campesterol** | 24α-methyl cholest-5-en-3b-ol | 5 | C1 | **0.585** | **0.005** | **0.240** | **0.011** |
| Campestanol | 24α-methyl cholestan-3b-ol | 0 | C1 | <0.001 | 1.000 | 0.290 | 0.119 |
| Avenasterol | 24-ethyl cholesta-7,24(28) trans-dien-3b-ol | 7 | C2 | <0.001 | 1.000 | 0.069 | 0.814 |
| **Schottenol** | 24α-ethyl cholest-7-en-3b-ol | 7 | C2 | **1.003** | **0.001** | **0.440** | **0.008** |
| **Sitosterol** | 24α-ethyl cholest-5-en-3b-ol | 5 | C2 | 0.275 | 1.000 | **0.183** | **0.012** |
| Sitostanol | 24α-ethyl cholestan-3b-ol | 0 | C2 | <0.001 | 1.000 | 0.093 | 0.606 |
| **Isofucosterol** | 24-ethyl cholesta-5,24(28) trans-dien-3b-ol | 5 | C2 | **0.634** | **0.002** | 0.141 | 0.053 |
| Stigmasterol | 24α-ethyl cholesta-5,22 trans-dien-3b-ol | 5 | C2 | <0.001 | 1.000 | 0.149 | 0.200 |
| **H-diversity** | Shannon Index |  |  | **0.834** | **0.001** | **0.177** | **0.004** |
| Grouped Sterols | Grouping description |  |  |  |  |  |  |
| **Sum_C0** | No substitution at C-24 |  |  | **0.906** | **0.001** | **0.196** | **0.025** |
| **Sum_C1** | CH<sub>n</sub> substitution at C-24 |  |  | **0.689** | **0.001** | **0.158** | **0.012** |
| **Sum_C2** | C<sub>2</sub>H<sub>n</sub> substitution at C-24 |  |  | **0.668** | **0.001** | **0.169** | **0.002** |
| **Sum_D0** | Phytosterols (Saturated ring B) |  |  | **0.880** | **0.004** | **0.177** | **0.032** |
| **Sum_D5** | $\Delta^5$ Sterols | | | **0.777** | **0.001** | **0.201** | **0.007** |
| **Sum_D7** | $\Delta^7$ Sterols | | | **0.825** | **0.001** | **0.188** | **0.035** |
| Sum_D8 | $\Delta^8$ Sterols | | | 0.022 | 0.114 | 0.167 | 0.152 |
| **Total sterol** |  |  |  | **1.002** | **0.001** | **0.293** | **0.026** |

### Sterols and abiotic environmental factors

How similar pollen sterol profiles are between plants was neither significantly associated with the similarity between native environmental niches (represented by environmental principal component axes PC1-PC3) nor with phylogenetic distances (r^2^ = 0.013, p = 0.17 for additive model in MRM analysis, for individual factors see Table 3).

**Table 3.** MRM (multiple regression on distance matrices) analysis results showing regression coefficients and p-values for the multiple regression of pairwise distances on the three first environmental principal components (PC1-PC3) and phylogenetic distances against the sterol profile Bray-Curtis distance matrix.

| Variable | Regression coefficient | p-value |
| --- | --- | --- |
| **Intercept** | 7.89E-01 | 0.24 |
| **Phylogenetic distance** | 2.27E-05 | 0.82 |
| **PC1** | 1.87E-02 | 0.0587 |
| **PC2** | -2.0 E-02 | 0.0585 |
| **PC3** | -1.07E-02 | 0.3635 |

Total pollen sterol content of plant species was positively correlated with some of the environmental variables in their native range, but in general the explained variance (r^2^) was low (Fig. 4, Table S4). Specifically, total sterol content correlated with environmental PC1 (associated with high mean temperatures, low temperature seasonality and low soil carbon content; p = 0.015, r^2^ = 0.060; Fig. 4). For linear models of individual environmental variables, species with higher total pollen sterol content tended to occur in locations with higher annual mean temperature, higher temperatures in the coldest quarter, higher precipitation in the wettest quarter, and lower temperature seasonality (p-values < 0.05 for linear models of phylogenetic independent contrasts, r^2^ between 0.05 to 0.08, Table S4), as is the case in tropical conditions. For Shannon’s H diversity of pollen sterol profiles, the only significant association with environmental variables was a weak negative correlation with temperature seasonality (p = 0.014; r^2^ = 0.06) (Table S4). None of the other environmental variables or principal components were significantly correlated with sterol content or diversity, nor was the total environmental niche breadth (Fig. 4, Table S4).

**Fig. 4.**
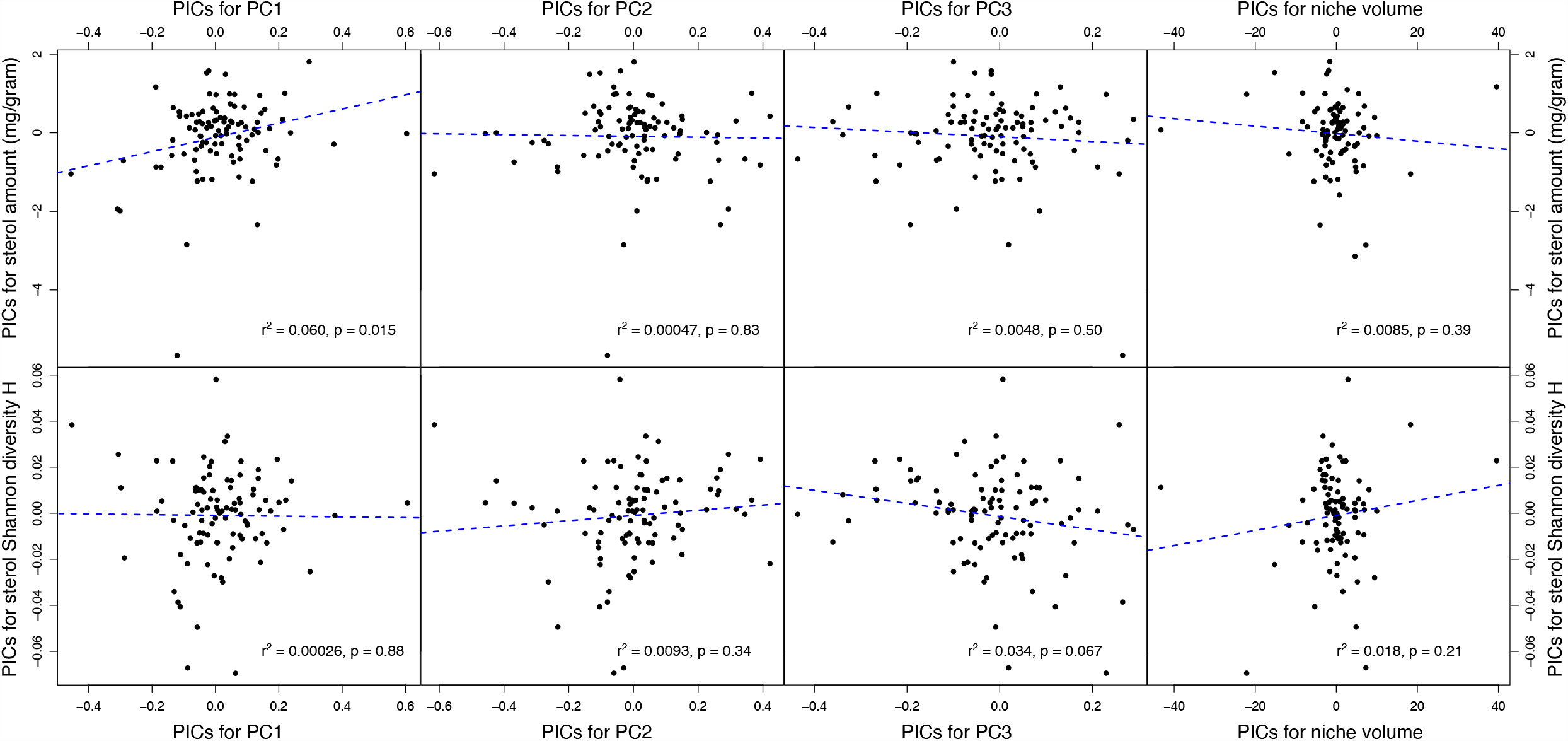
Correlation plots of phylogenetically independent contrasts (PICs) of positions on the environmental principal component axes (PC1-PC3) and environmental niche breadth against PICs of total pollen sterol amounts (top row) and sterol profile Shannon diversities H (bottom row). Blue dashed lines indicate regression lines of linear models (with intercept set to zero); r^2^ and p-values for linear models inserted in the respective plot. PC loadings from each environmental variable see Table S3.

### Sterols and pollinator guilds

We found overall pollen sterol profiles were largely overlapping between plant groups with different pollinator guilds (bee, Lepidoptera, generalist insect, bird, unknown; Fig. 5a; ANOSIM: among group R = -0.0069, p = 0.57; no significant difference for any pairwise group comparison), and between plants with or without pollen as reward for bee pollinators (Fig. 5b; ANOSIM: R = 0.033, p = 0.15). This suggests that pollinator guilds or the use of pollen as reward by bees do not explain differences in pollen sterol composition. We note that wind pollinated Angiosperms were not part of this dataset.

**Fig. 5.**
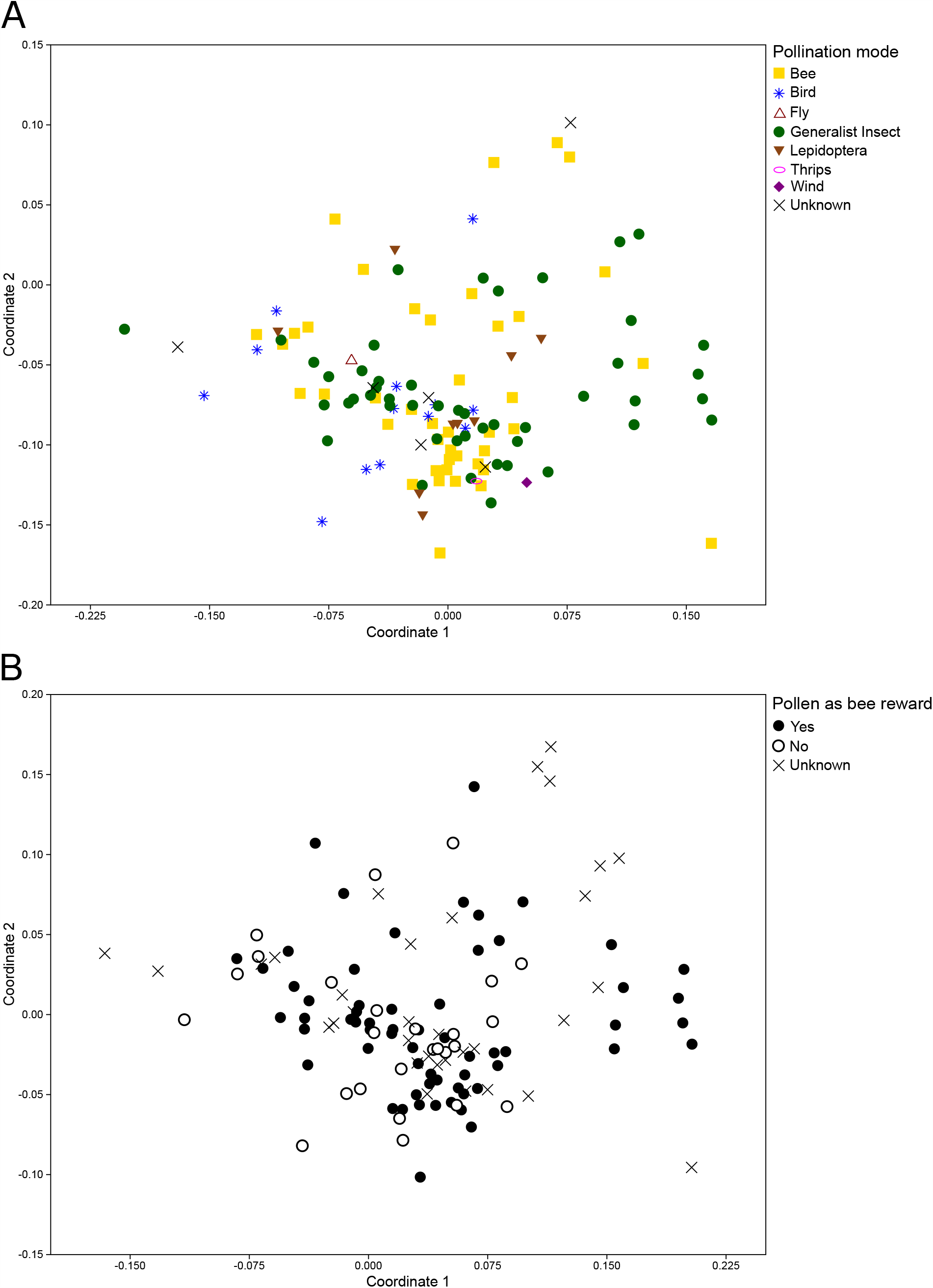
2D-NMDS plots of sterol profiles for plants (a) with different pollinator guilds, and (b) with/without pollen as bee reward. Distances correspond to sterol profile dissimilarity (Bray-Curtis distances). Stress of NMDS solution: 0.202.

Neither 24-methylenecholesterol nor total sterol content differed between plants with/without pollen as reward for bees (phylogenetic ANOVA: p = 0.46 and 0.66 respectively; Fig. S2, S3).

## Discussion

Phytosterols are primary metabolites in plants and are also essential nutrients for phytophagous insects, making them an important functional trait that provides a mechanistic link between plants and insects. Our study focused on the pollen sterol patterns across plant taxa, aiming to provide a more comprehensive overview of pollen sterol diversity and its relationship with plant phylogeny, abiotic environmental conditions, and pollinator guilds. We analysed 25 phytosterols in the pollen of more than 120 angiosperms representing 51 plant families and identified covariance patterns these phytosterols. Our data are the first to show significant phylogenetic signal for pollen phytosterols. Although environmental factors and pollinator guilds showed either weak or no relationships with pollen sterol content, future studies with more stratified sampling based on more finely defined pollinator species and environmental conditions could bring more insights about the drivers and importance of pollen sterol diversity.

### Phylogenetic structure of pollen sterols

Similar pollen sterol profiles in related taxa are ultimately due to shared evolutionary history and proximately due to shared genes for the enzymes involved in their biosynthesis. Indeed, we show that phylogenetic patterns in pollen sterols in part reflect their relations in the underlying biosynthesis pathway. For example, we observed significant phylogenetic clustering of plant species whose pollen sterol profiles are dominated by 24-methyl (C1-group), 24-ethyl (C2-group) or non-substituted (C0-group) phytosterols (Fig. 2, Table 2), reflecting the bifurcation of biosynthesis pathways (Fig. 3). Key enzymes (SMTs, sterol methyltransferases) that bifurcate the phytosterol pathways are SMT1, which methylates C0 sterol cycloartenol to 24-methylenecycloartanol, and SMT2, the key and effective enzyme to methylate 24-methyl to 24-ethyl sterols (Akihisa et al., 1991; Nes, 2000; Schaeffer *et al*., 2001; Neelakandan *et al*., 2009). Based on our findings, it would seem likely that the expression of these enzymes follows phylogenetically conserved patterns in different clades. Similarly, phylogenetic clusters of the main sterol groups based on the presence or absence of and position of the double bond in ring B (e.g., Δ^5^, Δ^7^, or saturated ring B) suggest conserved expressions of specific desaturases (e.g. STE1 or Δ^7^ and Δ^5^-sterol-C5-desaturases which convert saturated carbon bonds to methines) and reductase (e.g. DWF5, sterol-Δ^7^ and Δ^5^-reductase which reduce methines to saturated bonds) (Benveniste, 2004; Villette *et al*., 2015).

Our factor analysis (Table 1) further revealed an inverse relationship between the abundance of the major Δ^24,28^ sterol (24-methylenecholesterol) and the C24,28-saturated sterols: campesterol, sitosterol and stigmasterol. This suggests an overall trade-off of these two groups, the balance of which may be governed by DWF1 (sterol-Δ^24^-reductase) activity. Data from previous studies (Villette *et al*., 2015; Vanderplanck *et al*., 2020a) suggests a similar inverse correlation between 24-methylenecholesterol and the 24C-ethyl sterols, although this has not been explicitly stated. A high ratio of 24-methylenecholesterol to C24,28-saturated sterols is evident in Cactaceae, Droseraceae, Rosaceae, Onagraceae and Paeoniaceae. Conversely, C24,28-saturated sterols are more abundant than 24-methylenecholesterol in Ericaceae, Primulaceae, Salicaceae and Amaryllidaceae. These families are not closely related, suggesting convergent evolution of sterol composition. Overall, this indicates an interplay of environmental selection pressures for particular structural groups and phylogenetic constraints of sterol biosynthesis enzyme expression.

The composition of phytosterols appears to be tissue-dependent (Nes, 1990; Nes et al., 1993). For example, 24-methylenecholesterol has been identified as the main pollen sterol in many Cactaceae (Fig. 2, Table S1) but is not abundant in their photosynthetic tissue (Lusby *et al*., 1993; Standifer *et al*., 1968; Salt *et al*., 1987; Li 1996). The unique functional roles in pollen development when compared to the sporophyte may contribute to the distinct sterol profiles in pollen. We observed strong correlations among early, cyclopropyl sterol intermediates of the sterol pathway, particularly 9b,19-cyclopropyl sterols (Table 1). Cycloartenol, 31-norcycloartenol and 24-methylenecycloartanol are correlated with each other: 31-norcycloartenol and 24-methylenecycloartanol are both derived from cycloartenol. 24,25-Dehydropollinastanol and 31-norcycloartanol also show high correlation and both are derived from 31-norcycloartenol. Co-occurrence of cyclopropyl sterols suggests a reduction in CPI1 (cyclopropyl isomerase) activity and truncation of the sterol pathway, either within the pollen grain or in the surrounding tapetum cells from which pollen coat sterols are derived. 9b,19-Cyclopropyl sterols have been identified as key components of the pollen coat in *Brassica napus* (Villette *et al*., 2015; Wu *et al*., 1999). In addition, cycloeuclanol is the main sterol synthesised in the growing pollen tube of *Nicotiana tabacum* (Villette *et al*., 2015; Rotsch *et al*., 2017).

### Correlations of phytosterols with abiotic factors

The presence of different phytosterols could be evolutionary adaptations to environmental conditions. We detected a positive relationship between sterol content and temperature (particularly mean annual temperature and mean temperature of the coldest quarter), and a negative correlation with temperature seasonality, even though the overall association strength was low (Table S4). This indicates that plants found in cool and temperate climatic conditions were likely to have less pollen sterol than those found in areas of the world with warm climates with little seasonal fluctuations (e.g., tropical climates). The association between warmer climates and higher total amounts of pollen sterols may have evolved as protection against membrane heat stress, since the role of phytosterols in adaptation to high temperature stress is established (Dufourc, 2008, Narayanan et al. 2016). Phytosterols including campesterol, sitosterol and avenasterol degrade in stored grain more rapidly at higher temperatures (Wawrzyniak *et al*., 2019), so higher sterol content in warmer climates may avert the risk of their rapid breakdown and limited availability. Besides this, other pollen sterol characteristics (e.g., sterol diversity and the overall pollen sterol composition) were not notably associated with abiotic factors. Our sampling, however, was biased towards plants of temperate regions (the predominant species available to us for sampling). Limited sampling towards extremes of the environmental gradients may have reduced our scope to detect associations between abiotic factors and pollen sterol characteristics. Future work should therefore be targeted at sampling additional plant species of more extreme environments to fill this gap. Note that our species were sampled at glasshouses (e.g., tropical glasshouse, alpine glasshouse) or outdoors at Royal Botanic Gardens Kew and nearby areas (sampling details see Table S1) to get a first estimate of pollen sterol diversity across a broad range of species. Future in-depth studies on how abiotic conditions affect pollen sterol variation within-species deserve further attention to build a more complete overview of pollen sterol diversity at different taxonomic levels.

### Impact of sterol diversity on pollinators

Pollen sterol amount and composition did not differ significantly between bee pollinated and non-bee pollinated plant species. This could indicate that pollen sterols have generally not been under selection by bee pollinators although we acknowledge that our analysis combined all bee pollinated plants into one group. Therefore, it remains possible that pollen sterols play a role in finer scale interactions between different bee species of varying levels of pollen specialization and their host plants. We also note that, although we based our assessment of pollinator guilds on the best available literature data, the quality of evidence for the effective pollinators of the plants in our data set varied. This calls for further in-depth studies of the relationships between pollen sterols and pollinators, also including wind-pollinated Angiosperm taxa missing in this work as points of comparison to animal pollinated plants.

A major knowledge gap exists in understanding how important specific phytosterols are for bees, particularly wild bee species, since many of them are pollen specialists. Plants adapt nectar chemistry to suit the specific needs of pollinators (Vandelook *et al*., 2019) and could similarly alter nutritional chemistry of pollen to optimize its nutritional suitability for flower visitors. Bee pollinators require a dietary source of sterols (Wright *et al*., 2018) and for this they must use the phytosterols found in pollen. Therefore, determining how lipid components of pollen vary qualitatively and quantitatively across different plant taxa is important in understanding how nutritionally limiting landscapes might be for bees, especially where they are not botanically diverse. For example, honeybee colony growth benefits from 24-methylenecholesterol (Herbert *et al*., 1980). Thus, honeybees may be nutritionally limited in landscapes where floral resources do not provision 24-methylenecholesterol. Our data suggested that many Asteraceae (e.g., *Achillea ptarmica* L., *Tanacetum vulgare* L. *Achillea millefolium* L., *Jacobaea vulgaris* Gaertn., *Centaurea nigra* L. and *Cirsium vulgare* (Savi.) Ten) are rich in Δ^7^-sterols (Fig. 2, Table S1) and lack the common honeybee-favourable Δ^5^-sterols (e.g., 24-methylenecholesterol). Δ^7^-sterols are known to be toxic to non-specialist herbivores and can only be utilized by some insect species (Behmer & Nes, 2003; Lang *et al*., 2012). Thus, plant species that produce unusual phytosterols in pollen may produce these as defence against pollen herbivory, but some specialist bee species may have developed mechanisms overcoming this defense. Indeed, pollen foraging bees on Asteraceae plants are mostly specialized oligolectic bees, while polylectic bee species avoid the pollen despite the ubiquitous distribution of Asteraceae species and their substantial amount of pollen provision (known as the Asteraceae paradox, Müller & Kuhlmann, 2008). While the reasons for this Asteraceae paradox remain unresolved, the abundance of Δ^7^-sterols we found in the pollen of Asteraceae species could provide a potential explanation (see also Vanderplanck *et al*., 2018, 2020b).

## Acknowledgements

We thank Wenxu Zhou for his guidance in analysing sterols, Amy Kendal-Smith, Richard Moore and James Woodward for their help with pollen collection, Geoffrey Kite for support with the GC-MS, Ellen Baker for helping revise the manuscript. We also thank the Horticultural staff of the Royal Botanic Gardens, Kew for maintenance of the outstanding living collections used to provide pollen for this study. The project was funded by a BBSRC grant to GAW (BB/P007449/1) and PCS (BB/P005276/1) and a Peter Sowerby Grant to PCS and HK. PZ was supported to work with DN at Texas Tech on a BBSRC International Travel Award Scheme grant (BB/S004653/1) to PCS.

## Author contributions

PZ, HK, and PCS designed the research. PZ collected the pollen and extracted and analysed sterols. DN, IWF and DIF helped with sterol identification and quantification. OS collected phylogenetic information on studied species and conducted phylogenetic analyses. SP, CP, and IO collected species geographic and environmental information and performed abiotic niche analyses. HK and PZ collected pollinator records on studied species, and HK conducted analyses with environmental factors and pollinator guilds. EM and GW conducted factor analysis. WDN generated sterol biosynthesis pathways. PZ and PCS drafted the manuscript. HK, OS, SP, GAW and all other authors contributed in writing and revising the manuscript.

## Supplementary materials

### Figures

**Fig. S1**. 2D-NMDS plot: Pollen sterol profile similarities between species of different plant families.

**Fig. S2**. Boxplot: 24-MC content (µg/mg pollen) of plants with/without pollen as bee reward.

**Fig. S3**. Boxplot: Sterol content (µg/mg pollen) of plants with/without pollen as bee reward.

**Fig. S4**. GC-MS spectra of the 25 phytosterols identified in our study (after Tri-sil derivatisation, extraction details see Materials and methods section).

### Tables

**Table S1**. Data table (plant species, scores for different environmental variables/principal components, pollinator guilds, sterol composition (relative & absolute amounts)).

**Table S2**. Scientific name and family for all sampled species, along with suggested OTL synonyms (which were subsequently used) and taxon IDs; species excluded from the phylogeny are highlighted in grey; reason for exclusion due to issues in the data and/or the OTL taxonomy are indicated.

**Table S3**. Variable contributions to axes of PCA of 13 environmental variables.

**Table S4**. Test results: Linear models of phylogenetic independent contrasts (PICs) of total sterol amount/diversity against PICs of environmental variables and niche breadth.

