## Supplementary for "Pollen sterols are associated with phylogenetics and environment but not with pollinators"

*New Phytologist* **Supporting Information**

**Article title:**

**Article acceptance date:** TBD

The following Supporting Information is available for this article:

**Supplementary figures:**

**Fig. S1.** 2D-NMDS (non-metric multudimensional scaling) plot of pollen sterol profile similarities (Bray-Curtis) between plant species with major plant families highlighted with different symbols (see legend). Distances between points correspond to sterol profile dissimilarities.

**Fig. S2.** 24-methylenecholesterol content (µg/mg pollen) of plants without pollen as bee reward (no bee pollination/collection of pollen by bees), or with pollen as reward for bees (based on evidence of bee pollination and pollen collection by bees).

**Fig. S3.** Total sterol content (µg/mg pollen) of plants without pollen as bee reward, or with pollen as reward for bees. Indentations represent 95% confidence intervals.

**Fig. S4.** GC-MS spectra of the 25 phytosterols identified in our study (after Tri-sil derivatisation, extraction details see Materials and methods section).

**Supplementary tables** (submitted separately: multiple sheets in Excel):

**Table S1.** Data table (plant species, scores for different environmental variables/principal components, pollination modes, sterol composition (relative & absolute amounts).

**Table S2.** Scientific name and family for all sampled species, along with suggested OTL synonyms (which were subsequently used) and taxon IDs; species excluded from the phylogeny are highlighted in grey; reason for exclusion due to issues in the data and/or the OTL taxonomy are indicated.

**Table S3.** Variable contributions to axes of PCA of 13 environmental variables.

**Table S4.** Results of linear model tests for phylogenetic independent contrasts (PICs) of total sterol amount/diversity against PICs of environmental variables and niche volume.

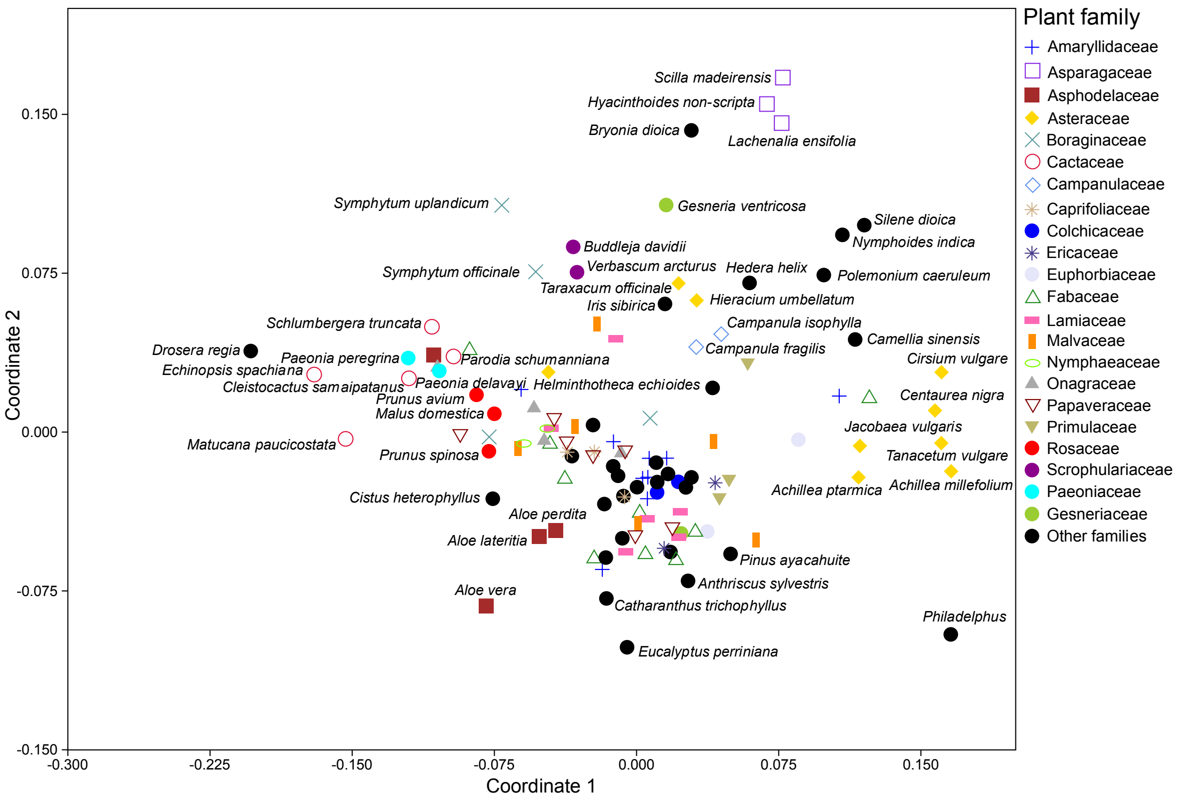


**Fig. S1.** 2D-NMDS (non-metric multidimensional scaling) plot of pollen sterol profile similarities (Bray-Curtis) between plant species with major plant families highlighted with different symbols (see legend). Distances between points correspond to sterol profile dissimilarities.

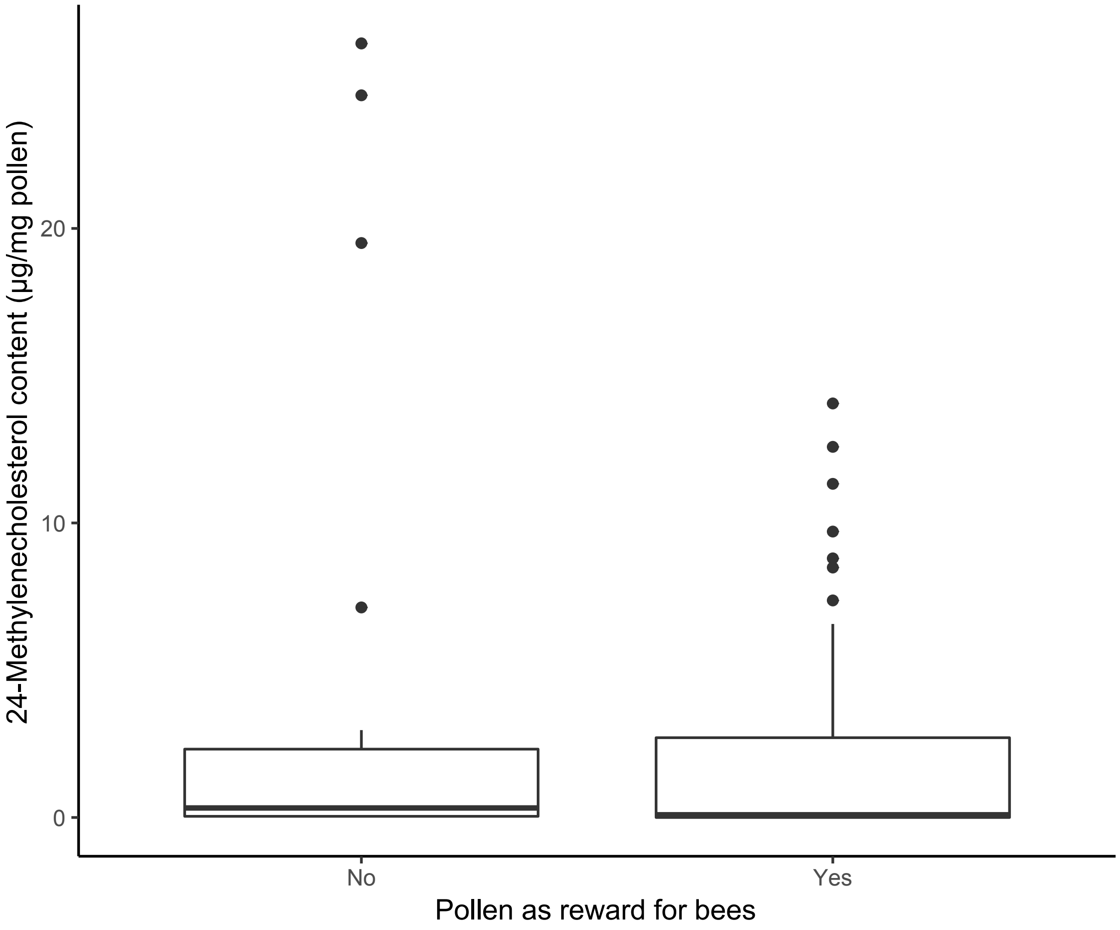


**Fig. S2.** 24-methylenecholesterol content (µg/mg pollen) of plants without pollen as bee reward (no bee pollination/collection of pollen by bees; n = 22), or with pollen as reward for bees (based on evidence of bee pollination and pollen collection by bees; n = 54).

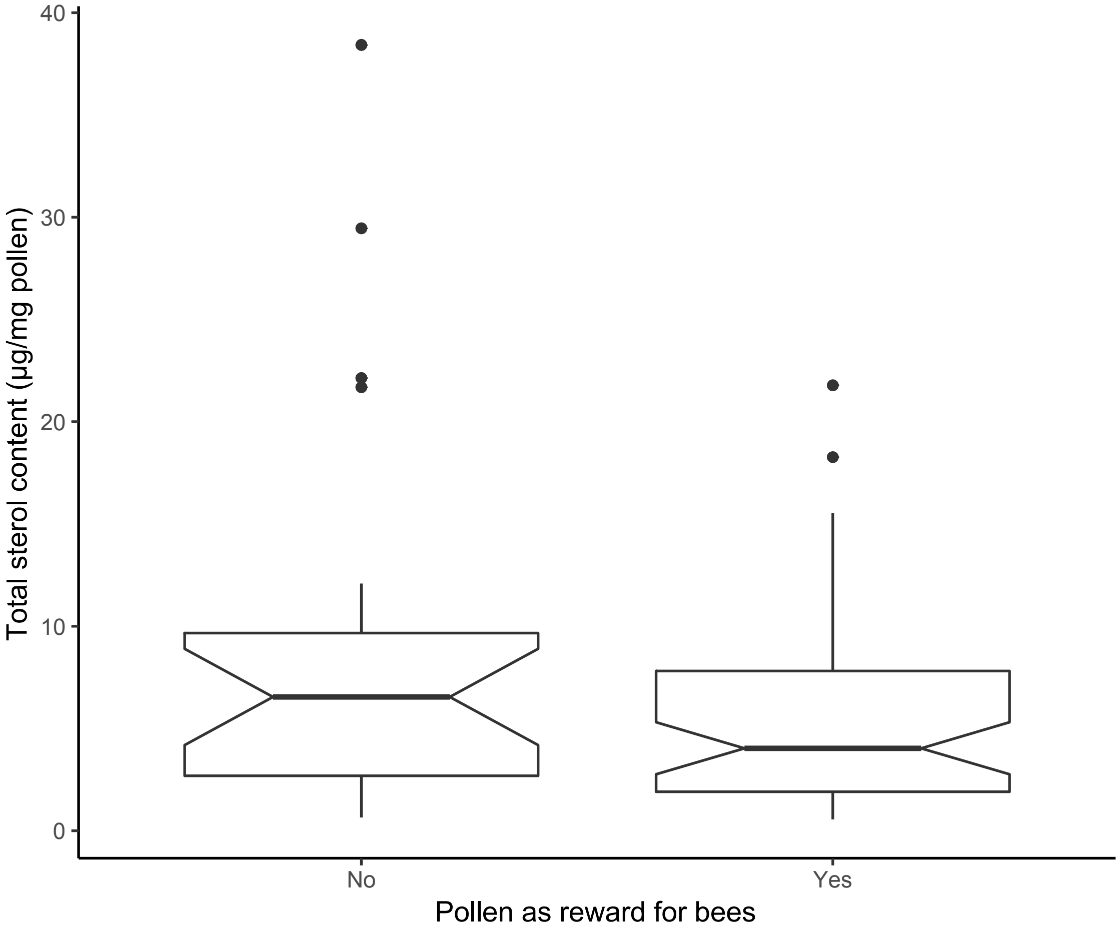


**Fig. S3.** Total sterol content (µg/mg pollen) of plants without pollen as bee reward (n = 22), or with pollen as reward for bees (n = 54). Indentations represent 95% confidence intervals.

**Fig. S4.** GC-MS spectra of the 25 phytosterols identified in our study (after Tri-sil derivatisation, extraction details see Materials and methods section).

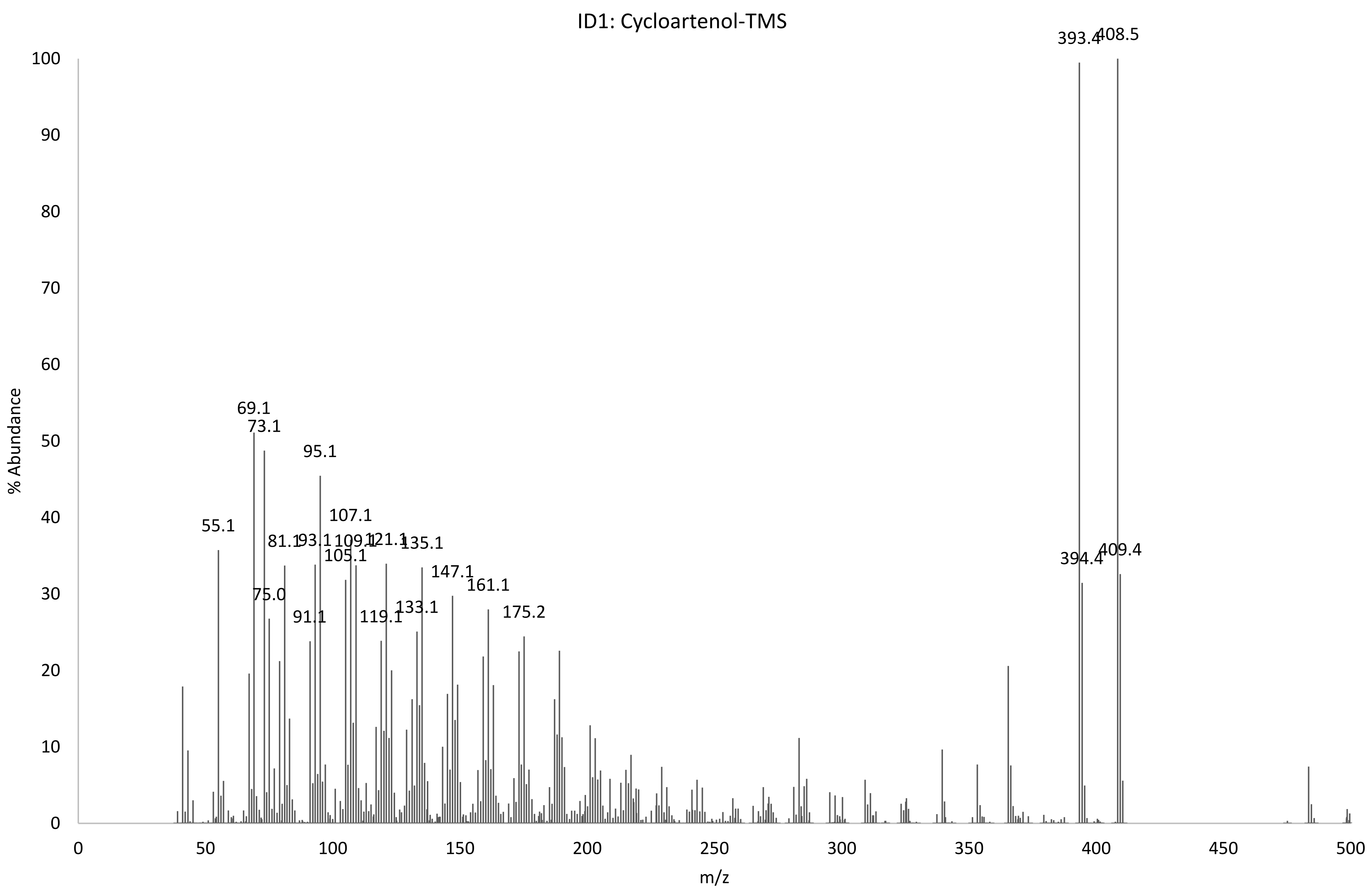
ID1: Cycloartenol-TMS
% Abundance
m/z
0
10
20
30
40
50
60
70
80
90
100
0
50
100
150
200
250
300
350
400
450
500
55.1
69.1
73.1
75.0
81.1
91.1
93.1
95.1
105.1
107.1
109.1
119.1
121.1
133.1
135.1
147.1
161.1
175.2
393.4
394.4
408.5
409.4

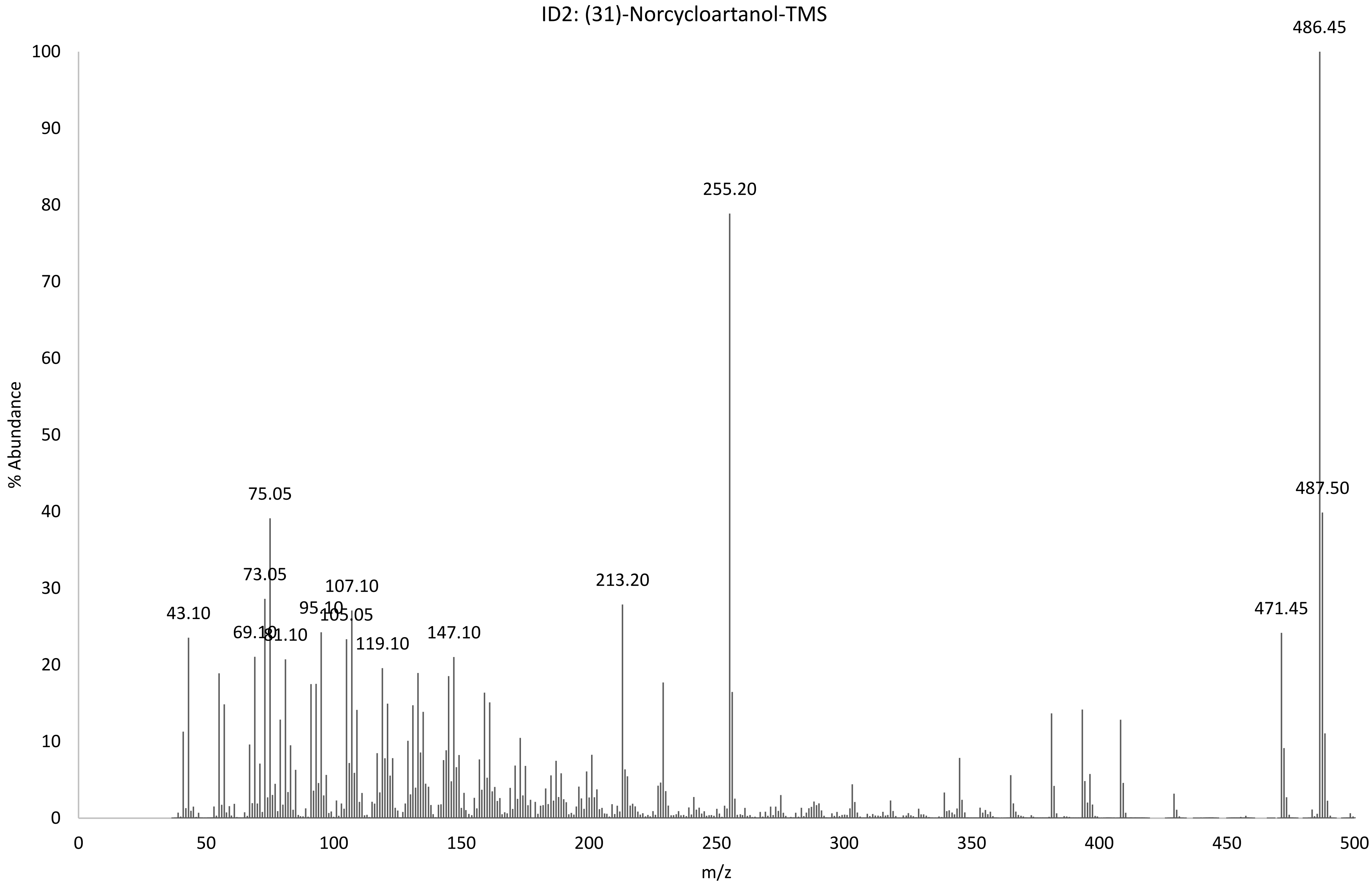
ID2: (31)-Norcycloartanol-TMS
% Abundance
m/z
486.45
487.50
471.45
255.20
213.20
147.10
119.10
107.10
105.05
95.10
75.05
73.05
81.10
43.10

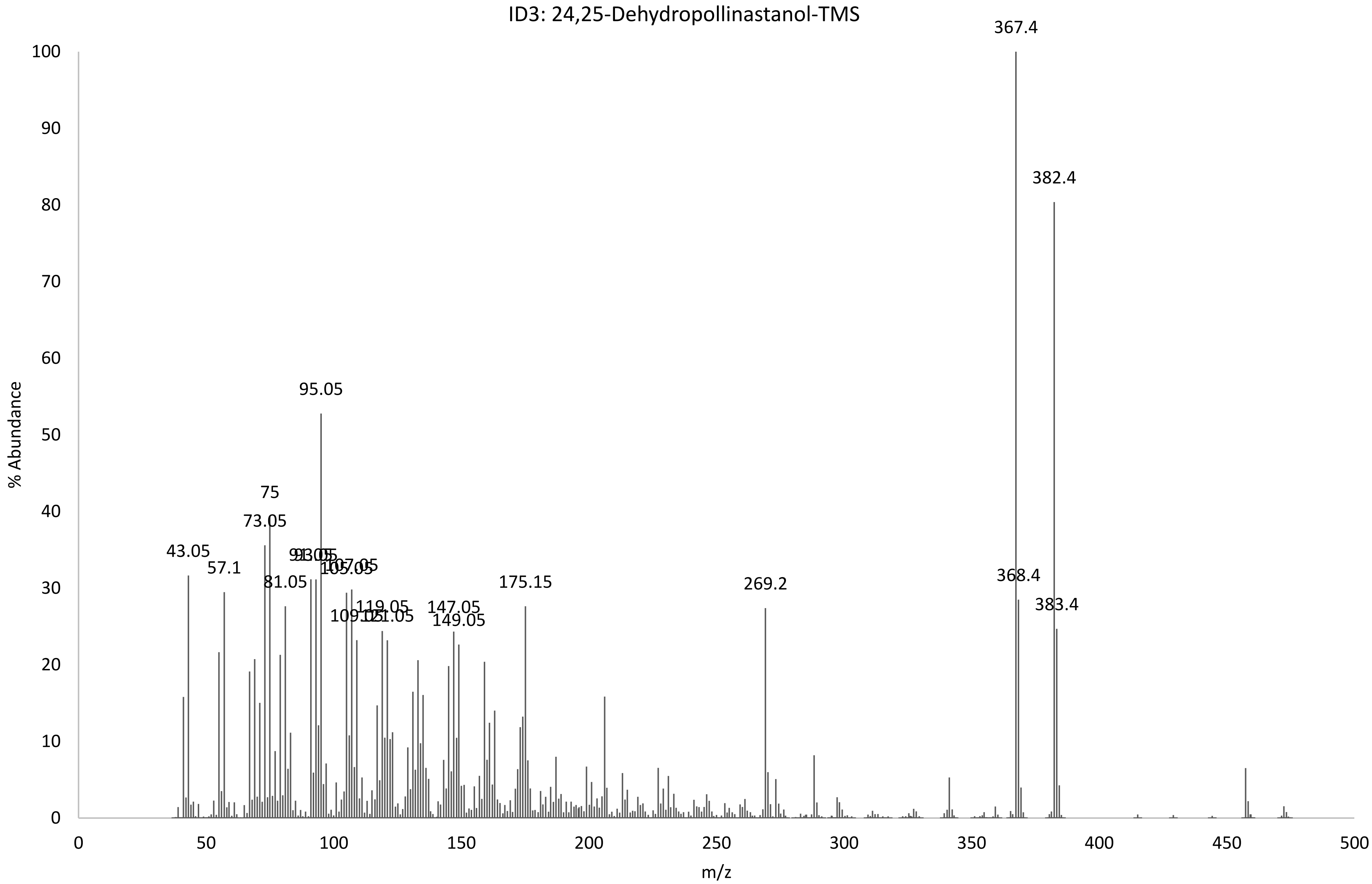
ID3: 24,25-Dehydropollinastanol-TMS
% Abundance
m/z
0
10
20
30
40
50
60
70
80
90
100
0
50
100
150
200
250
300
350
400
450
500
43.05
57.1
73.05
75
81.05
95.05
105.05
107.05
119.05
147.05
149.05
175.15
269.2
367.4
368.4
382.4
383.4

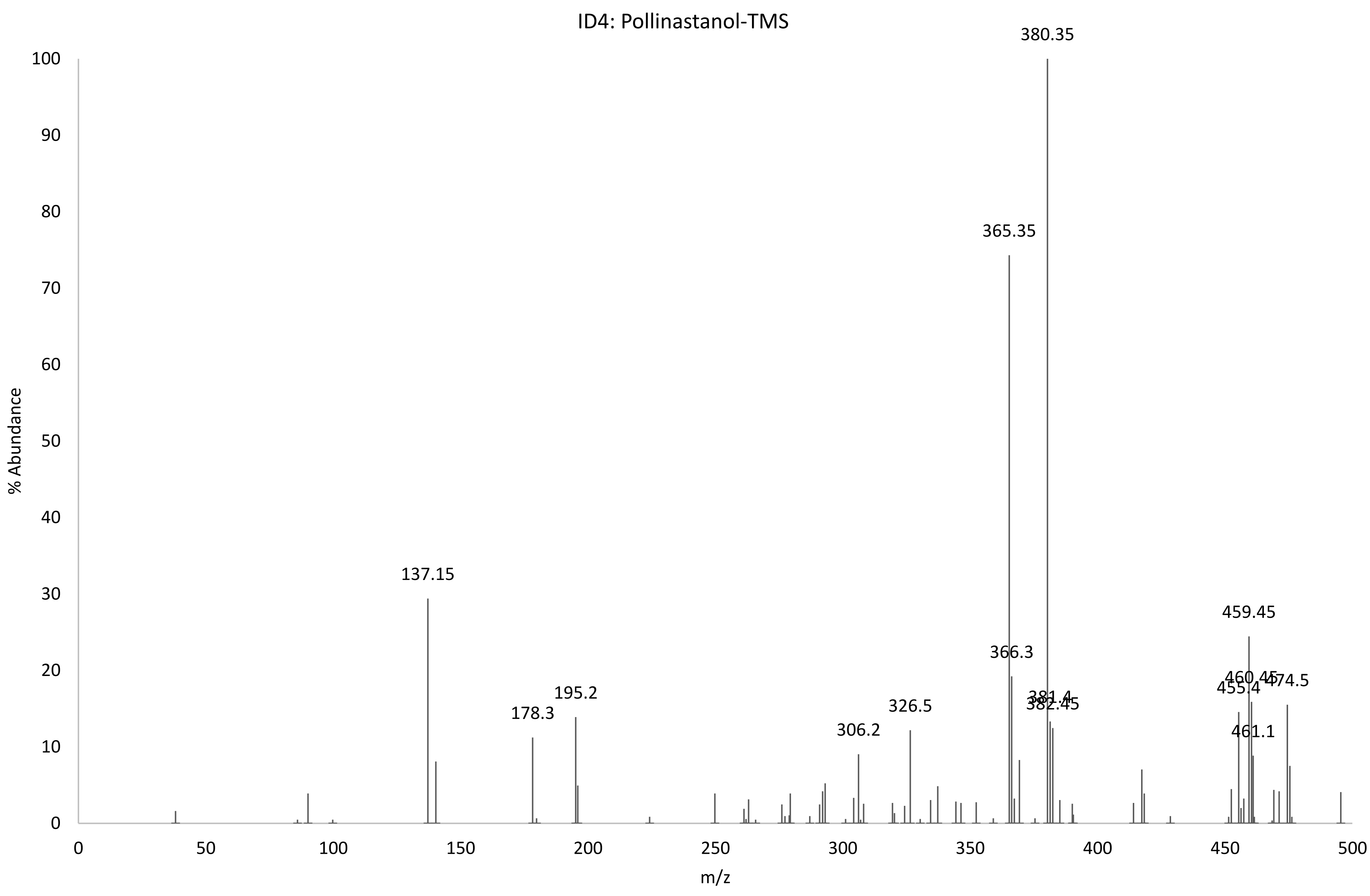

ID4: Pollinastanol-TMS
% Abundance
m/z
380.35
365.35
137.15
459.45
366.3
195.2
178.3
326.5
306.2
381.4
382.45
455.4
460.45
474.5
461.1
0
10
20
30
40
50
60
70
80
90
100
0
50
100
150
200
250
300
350
400
450
500

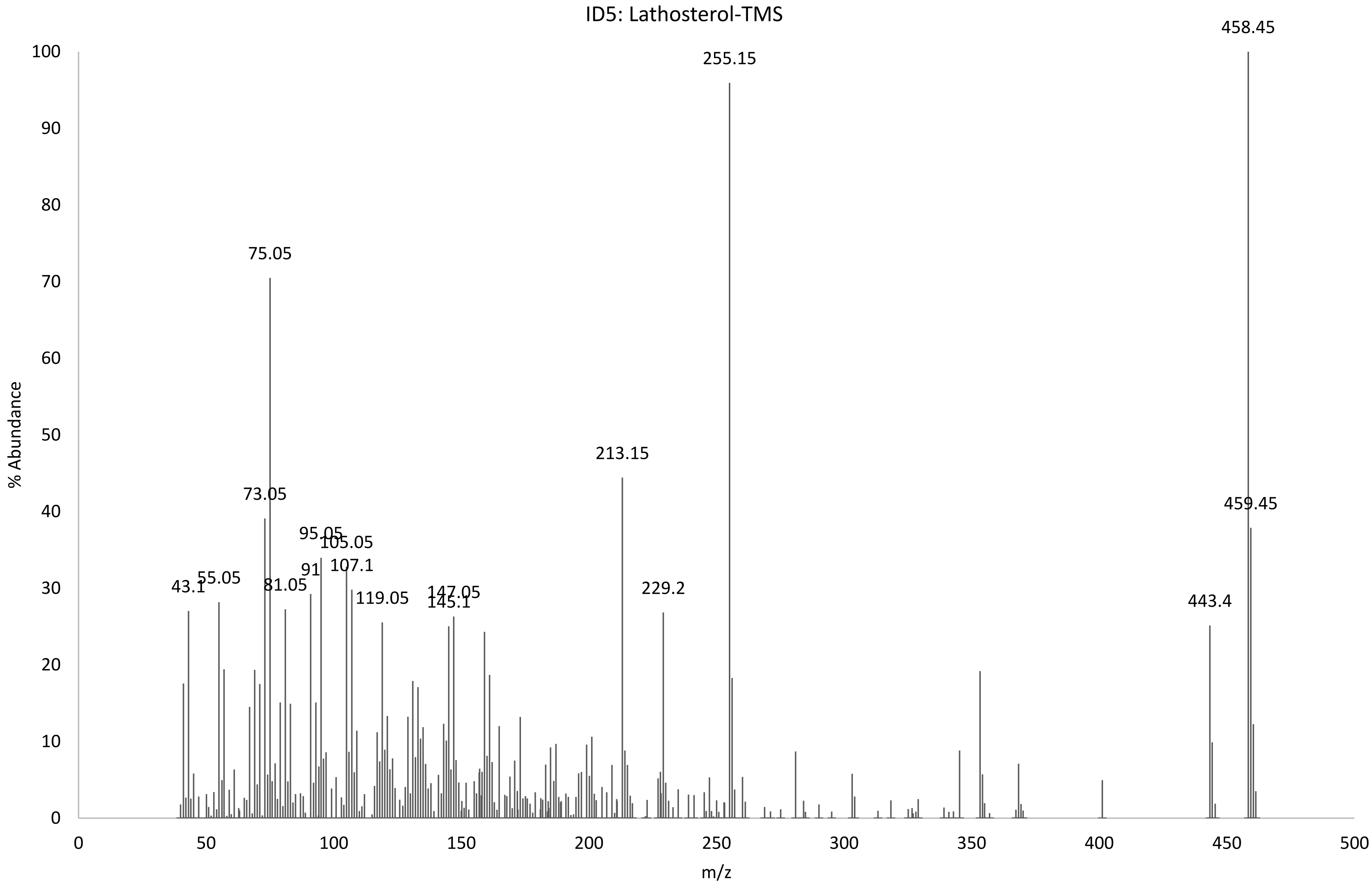
ID5: Lathosterol-TMS
% Abundance
m/z
0
10
20
30
40
50
60
70
80
90
100
0
50
100
150
200
250
300
350
400
450
500
43.1
55.05
73.05
75.05
81.05
91
95.05
105.05
107.1
119.05
145.1
147.05
213.15
229.2
255.15
443.4
458.45
459.45

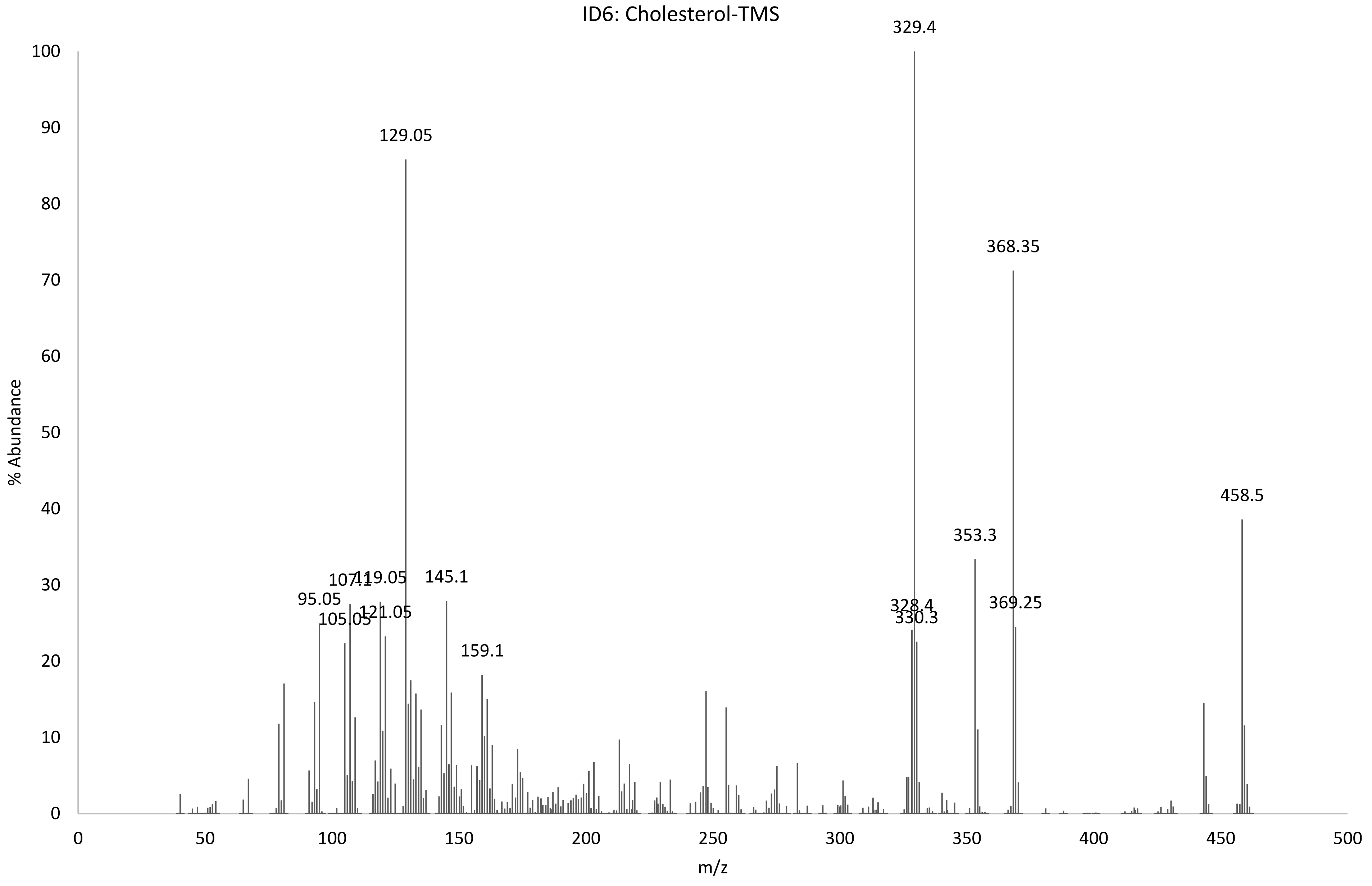
ID6: Cholesterol-TMS
% Abundance
m/z
0
10
20
30
40
50
60
70
80
90
100
0
50
100
150
200
250
300
350
400
450
500
129.05
95.05
105.05
107.1
119.05
121.05
145.1
159.1
329.4
328.4
330.3
353.3
368.35
369.25
458.5

ID7: 31-Norcyclartenol-TMS

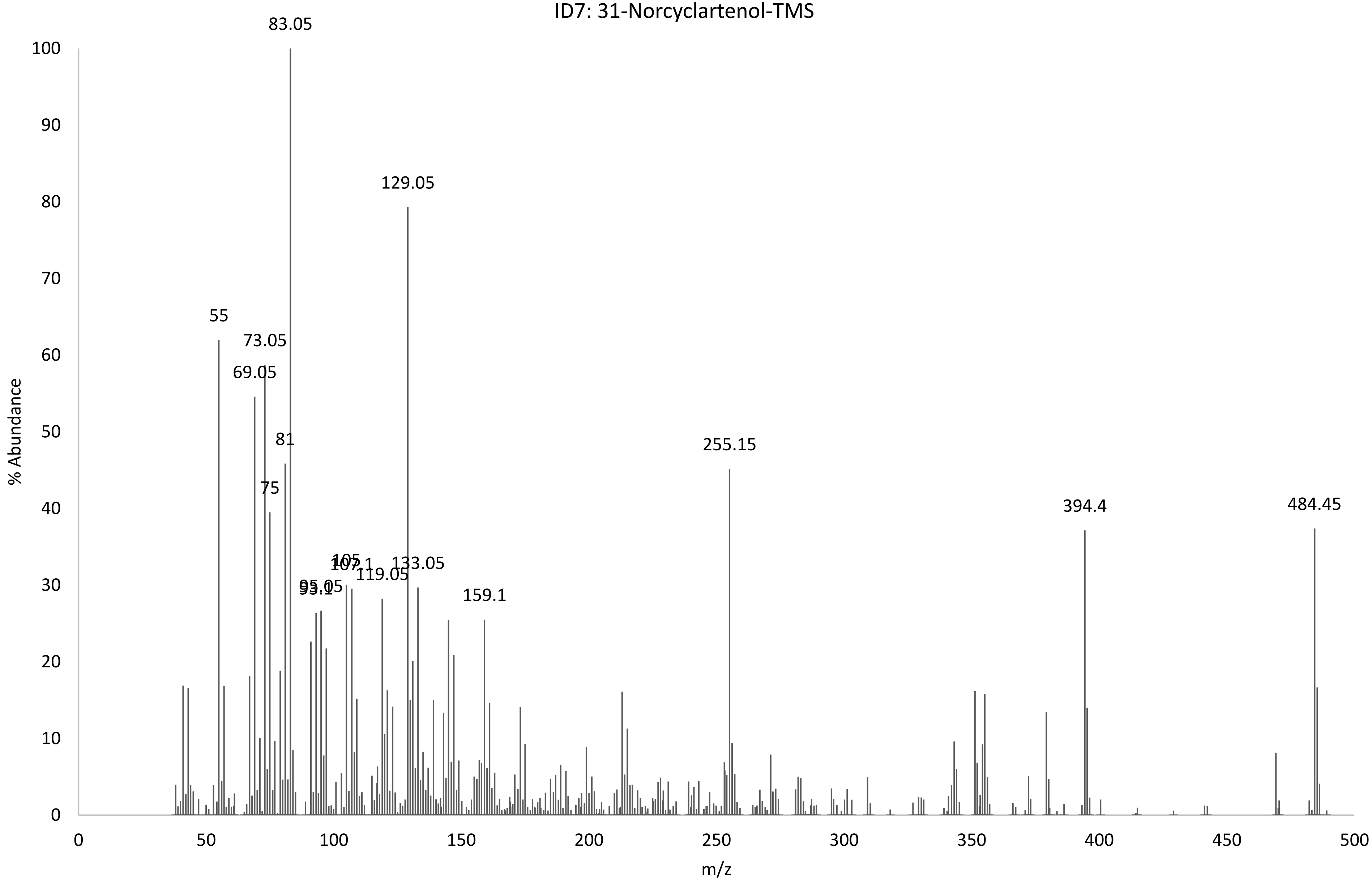

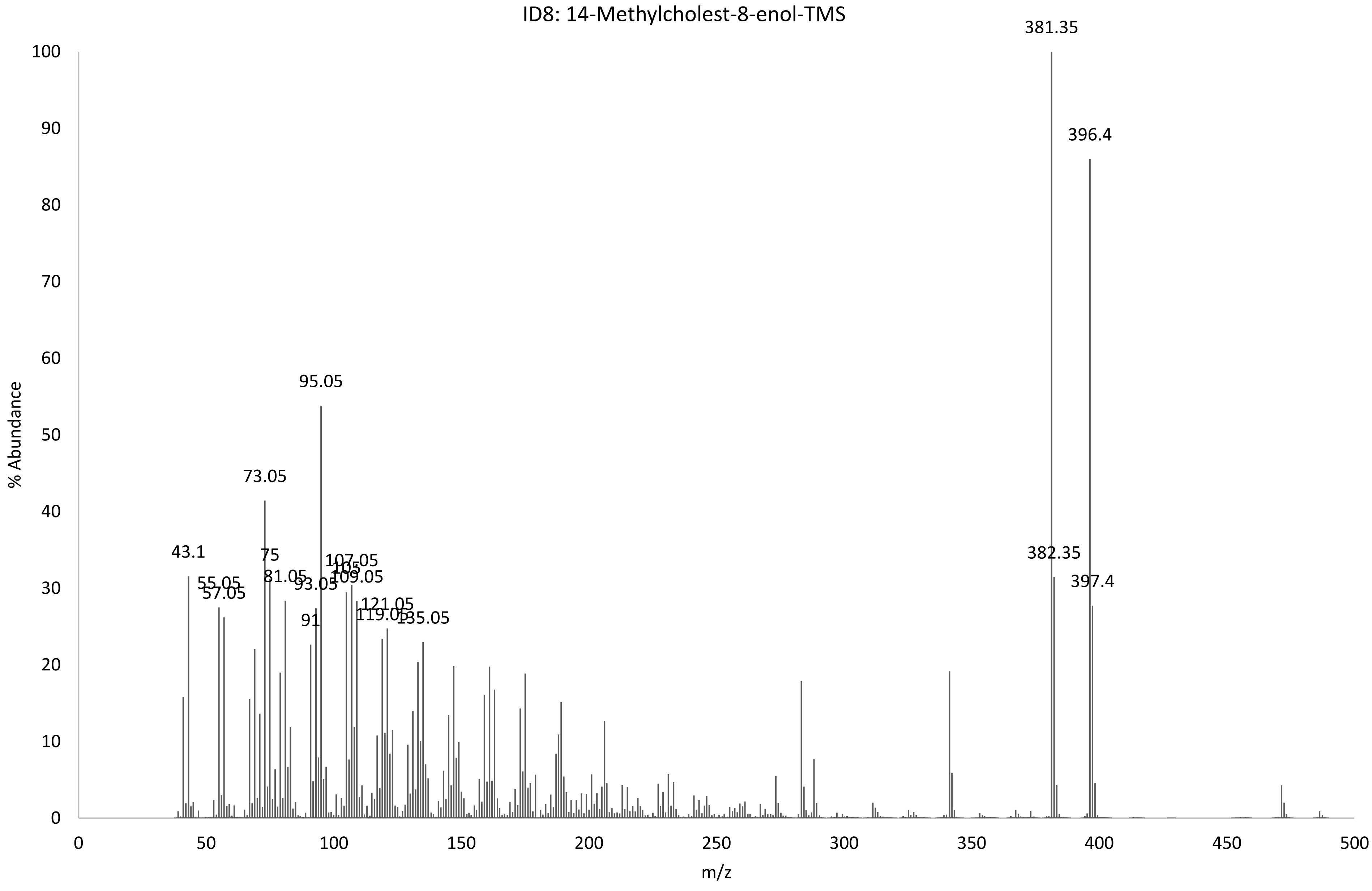
ID8: 14-Methylcholest-8-enol-TMS
% Abundance
m/z
381.35
396.4
95.05
73.05
43.1
75
107.05
105
109.05
55.05
57.05
81.05
93.05
121.05
119.05
135.05
91
382.35
397.4

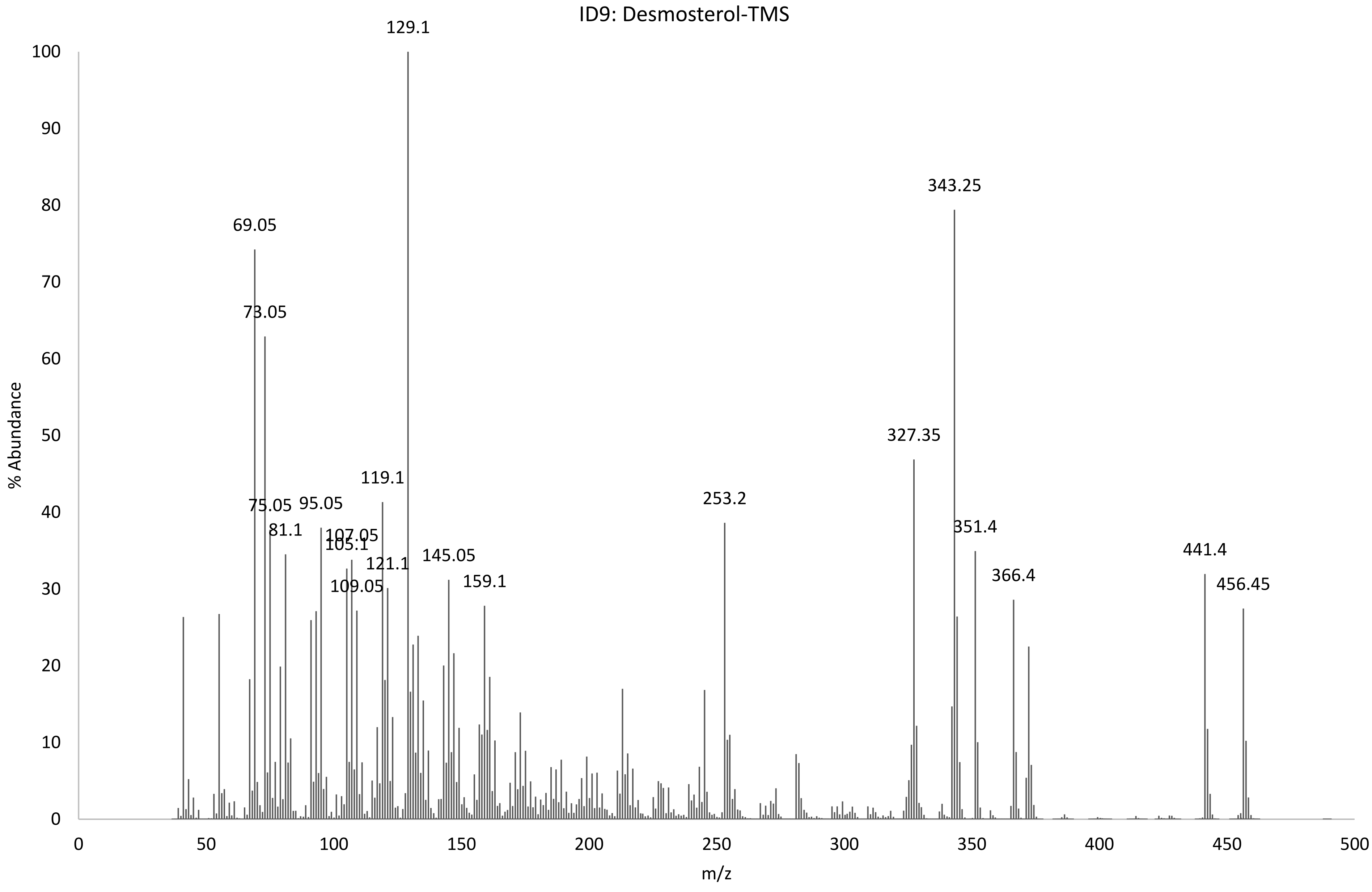
ID9: Desmosterol-TMS
% Abundance
m/z
129.1
343.25
69.05
73.05
327.35
119.1
75.05
95.05
253.2
81.1
107.05
105.1
351.4
441.4
145.05
121.1
366.4
109.05
159.1
456.45

ID10: 24-Methylenecholesterol-TMS

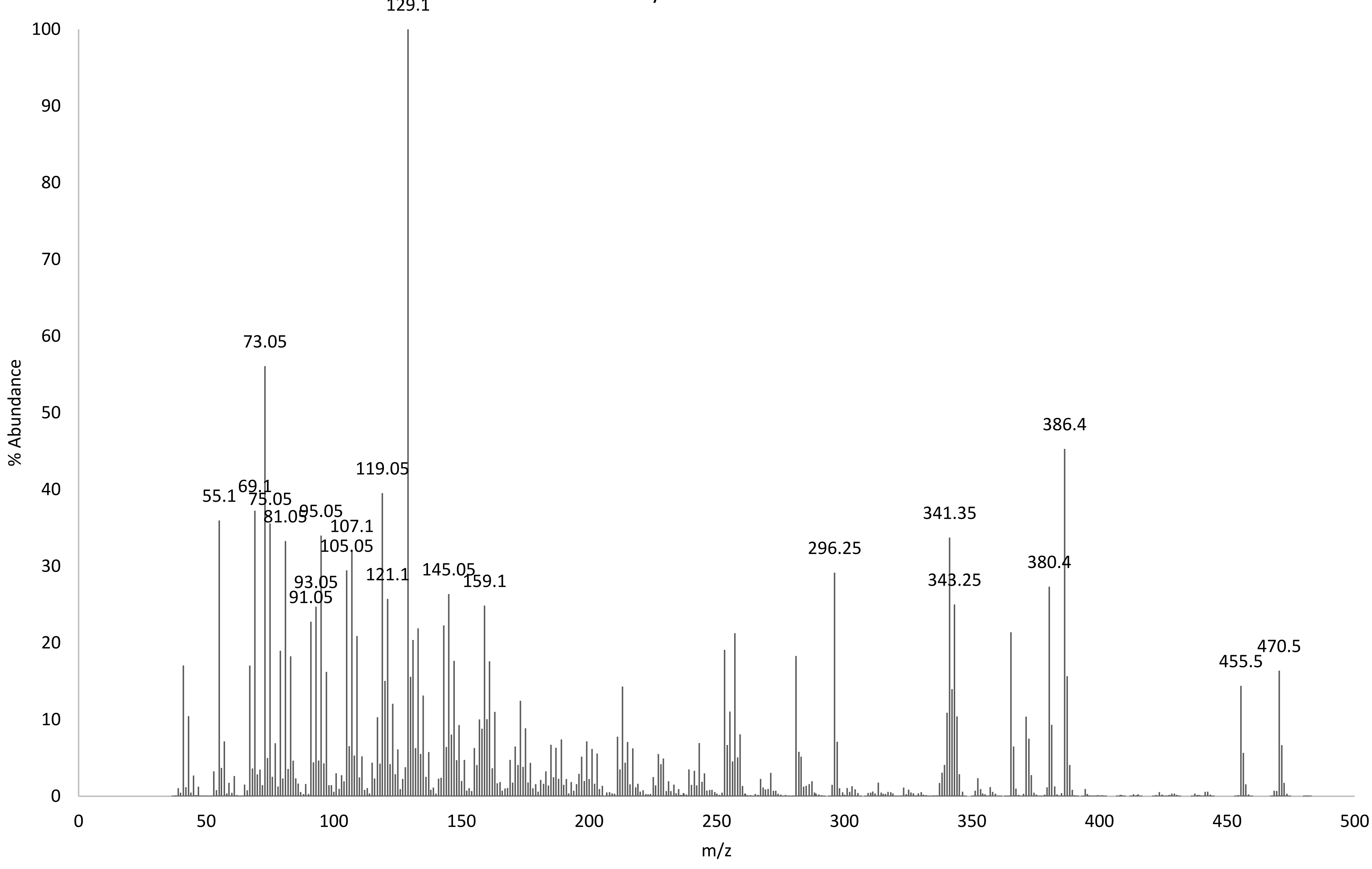

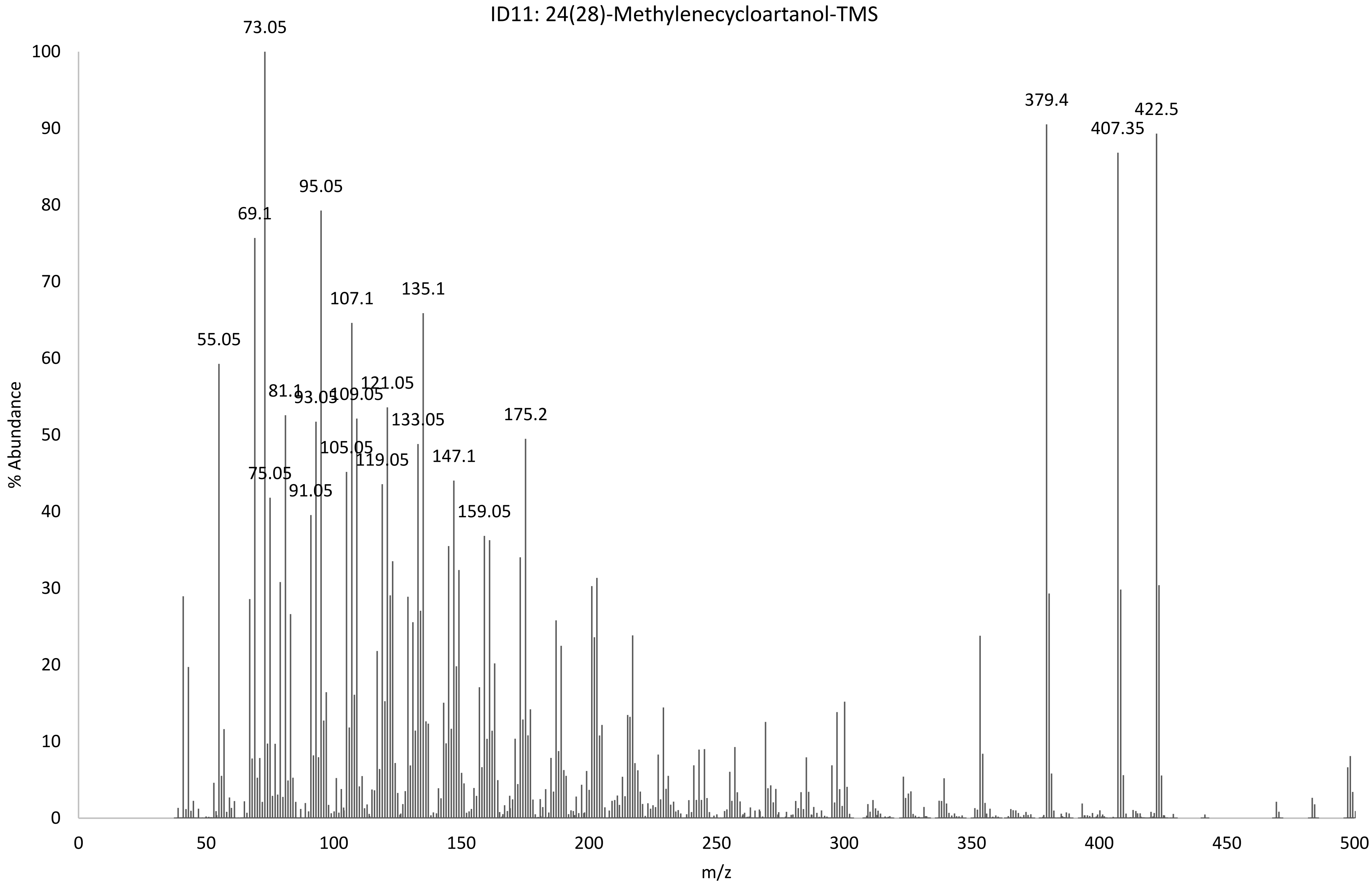
ID11: 24(28)-Methylenecycloartanol-TMS
73.05
95.05
69.1
107.1
135.1
55.05
121.05
81.1
93.05
109.05
133.05
175.2
105.05
119.05
147.1
75.05
91.05
159.05
379.4
407.35
422.5
% Abundance
m/z
0
10
20
30
40
50
60
70
80
90
100
0
50
100
150
200
250
300
350
400
450
500

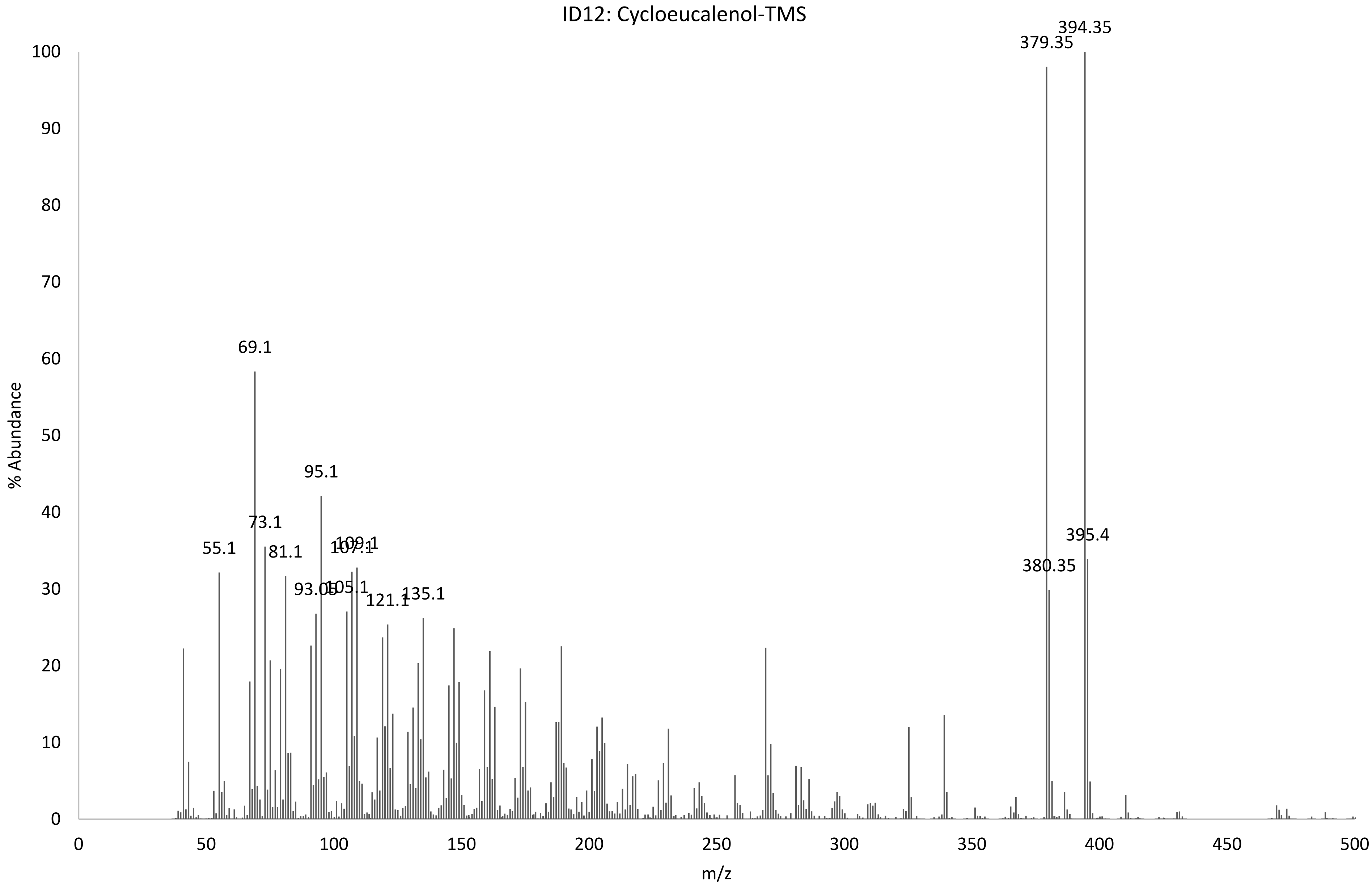
ID12: Cycloeucalenol-TMS
% Abundance
m/z
0
10
20
30
40
50
60
70
80
90
100
0
50
100
150
200
250
300
350
400
450
500
55.1
69.1
73.1
81.1
93.05
95.1
105.1
107.1
109.1
121.1
135.1
379.35
380.35
394.35
395.4

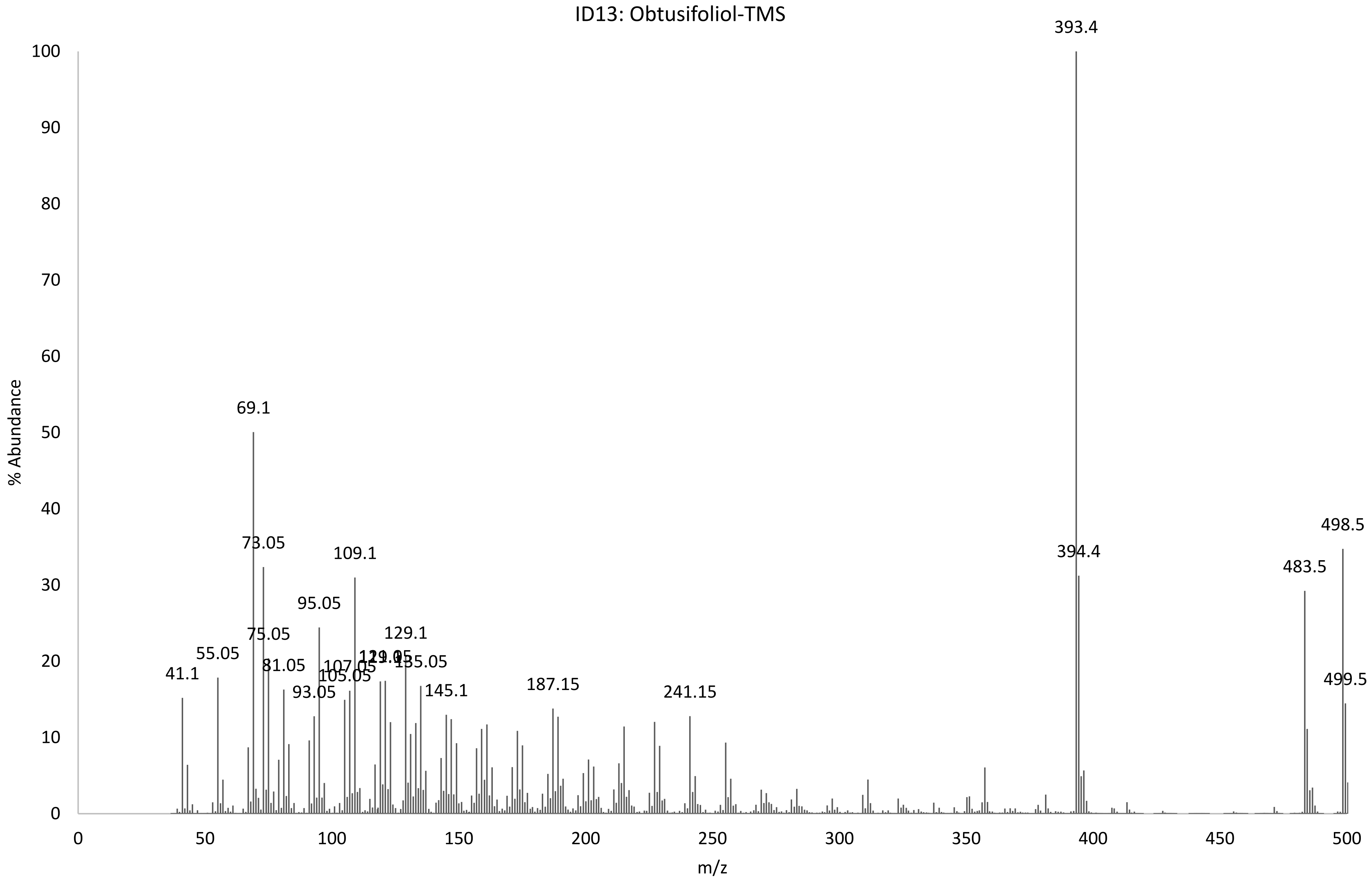
ID13: Obtusifoliol-TMS
% Abundance
m/z
393.4
69.1
73.05
109.1
95.05
75.05
129.1
55.05
81.05
107.05
135.05
41.1
105.05
93.05
145.1
187.15
241.15
394.4
483.5
498.5
499.5
0
10
20
30
40
50
60
70
80
90
100
0
50
100
150
200
250
300
350
400
450
500

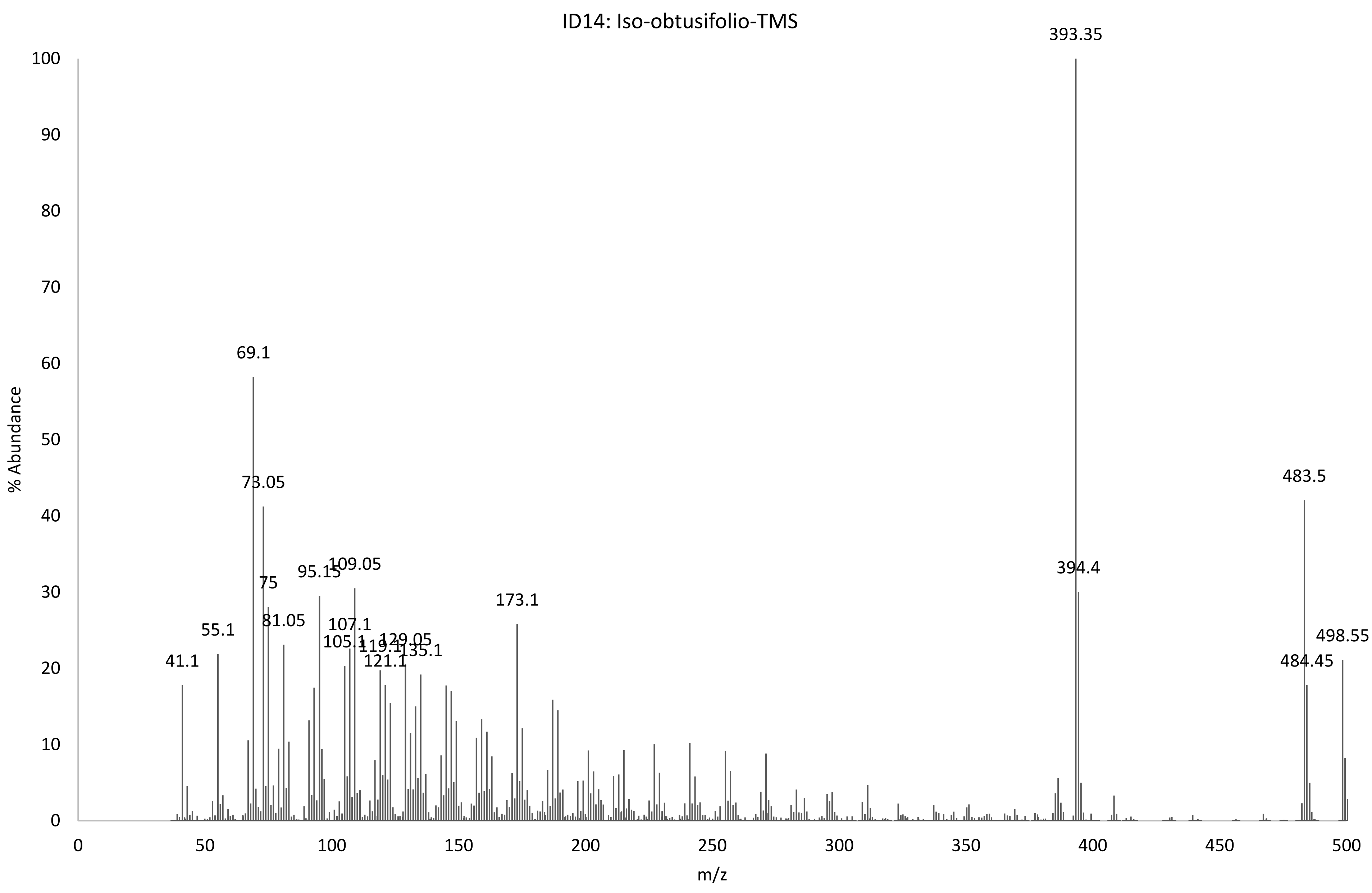
ID14: Iso-obtusifolio-TMS
% Abundance
m/z
393.35
394.4
483.5
484.45
498.55
69.1
73.05
75
81.05
55.1
41.1
95.15
109.05
107.1
105.1
119.1
129.05
121.1
135.1
173.1
0
10
20
30
40
50
60
70
80
90
100
0
50
100
150
200
250
300
350
400
450
500

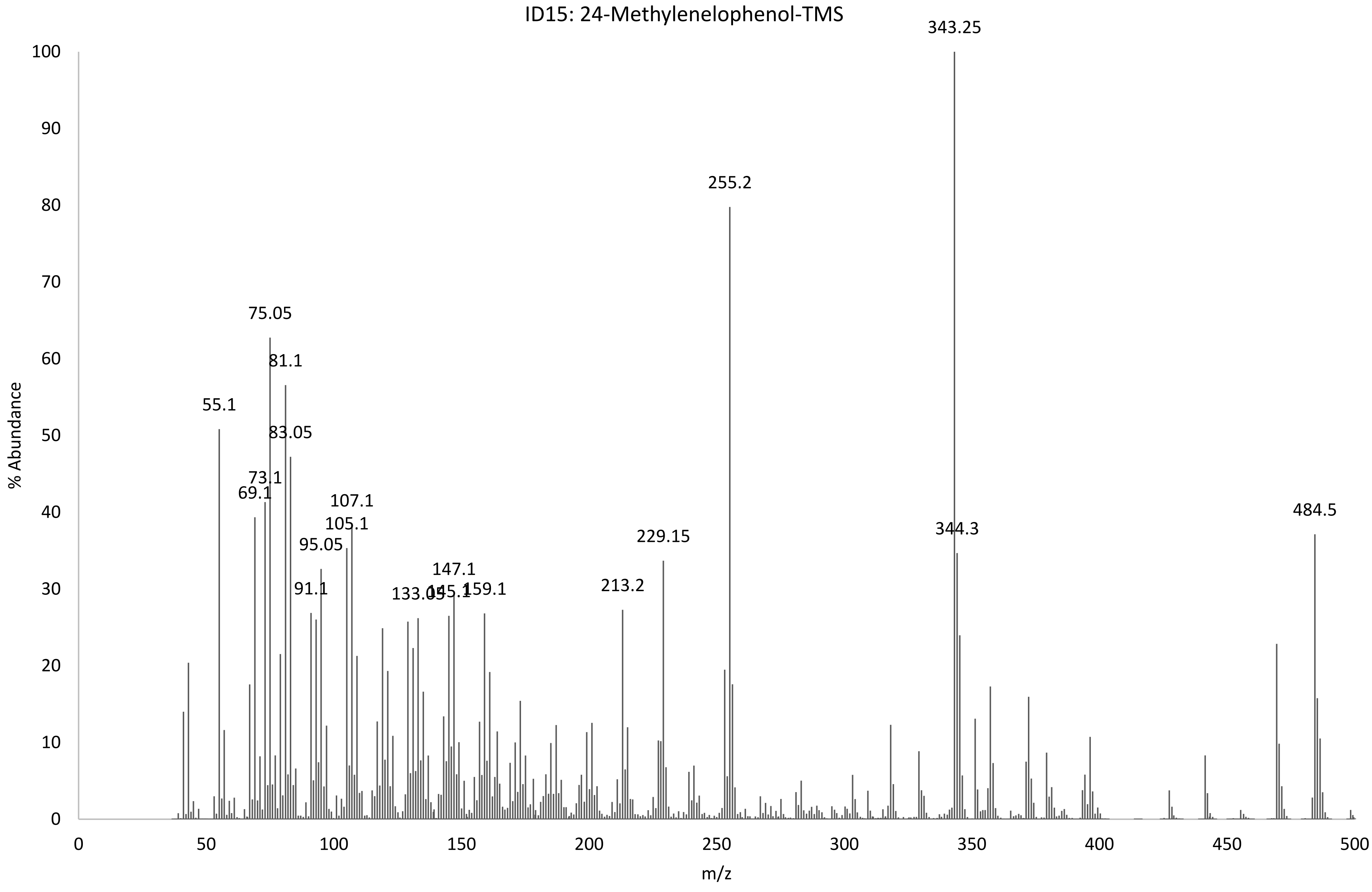

ID15: 24-Methylenelophenol-TMS
% Abundance
m/z
0
10
20
30
40
50
60
70
80
90
100
0
50
100
150
200
250
300
350
400
450
500
55.1
69.1
73.1
75.05
81.1
83.05
91.1
95.05
105.1
107.1
133.05
145.1
147.1
159.1
213.2
229.15
255.2
343.25
344.3
484.5

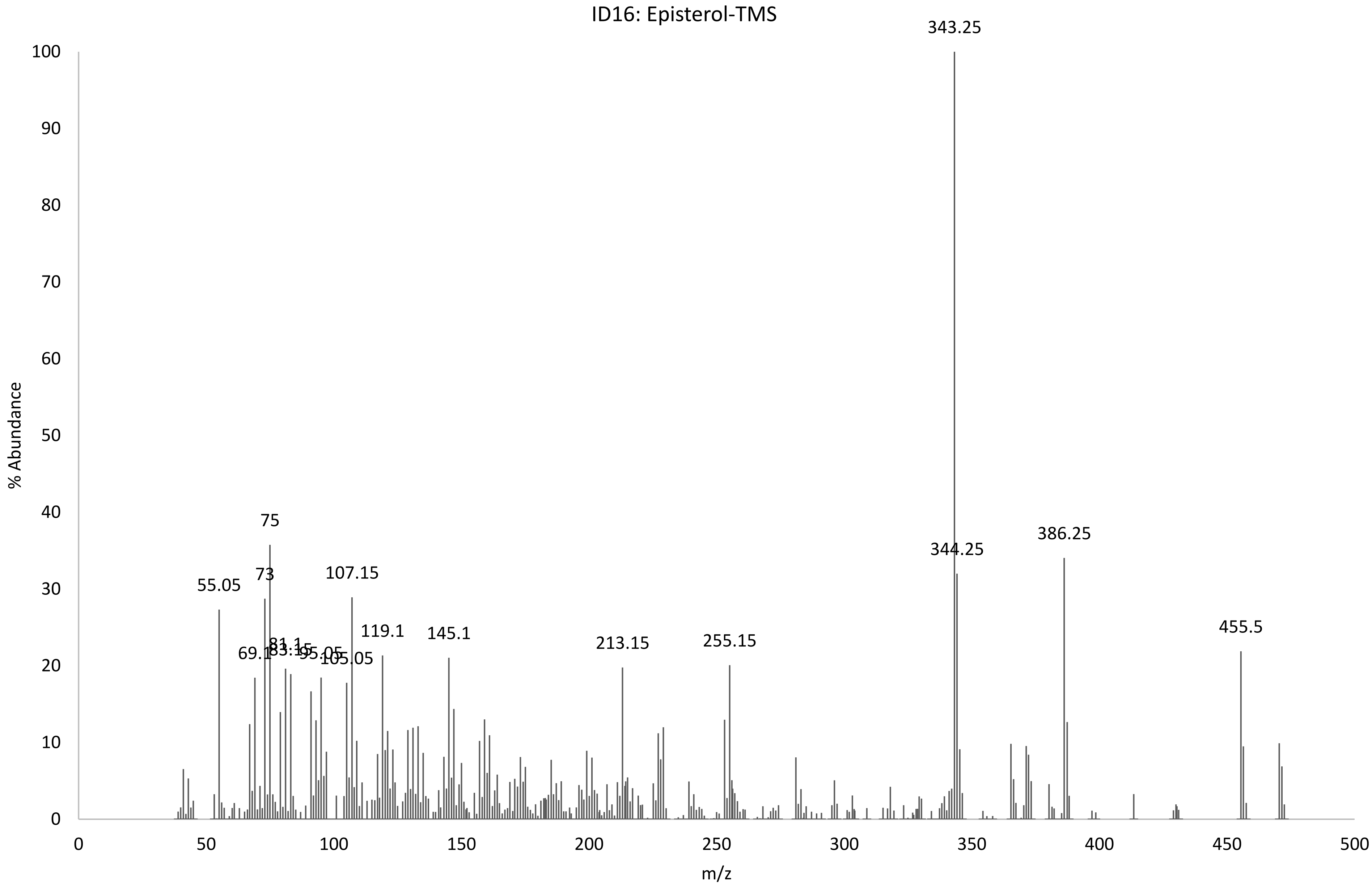

ID16: Episterol-TMS
343.25
344.25
386.25
455.5
75
73
55.05
107.15
119.1
145.1
213.15
255.15
69.1
81.1
83.15
95.05
105.05
% Abundance
m/z
0
10
20
30
40
50
60
70
80
90
100
0
50
100
150
200
250
300
350
400
450
500

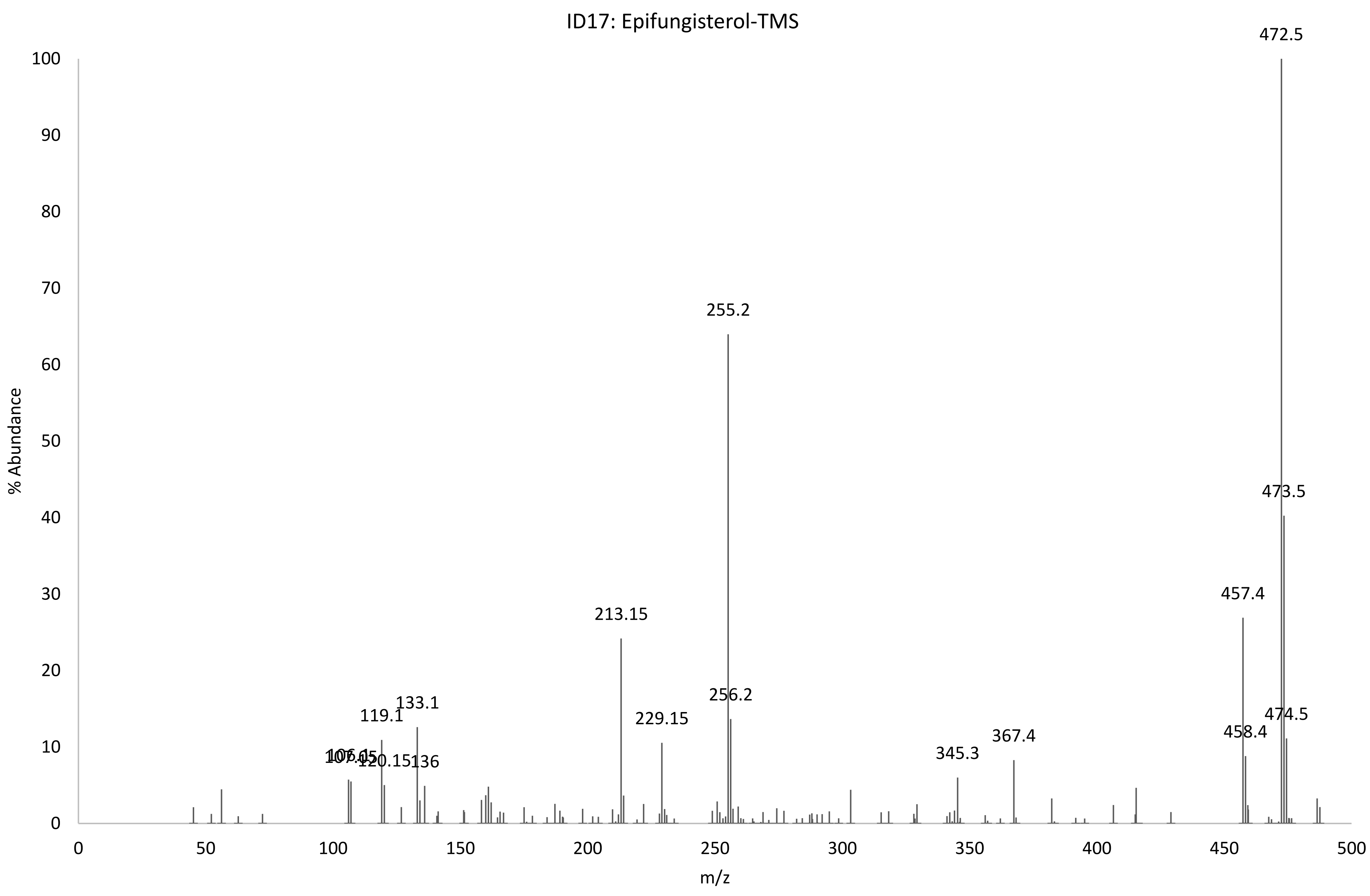

ID17: Epifungisterol-TMS
% Abundance
m/z
472.5
473.5
474.5
457.4
458.4
367.4
345.3
255.2
256.2
229.15
213.15
133.1
119.1
136

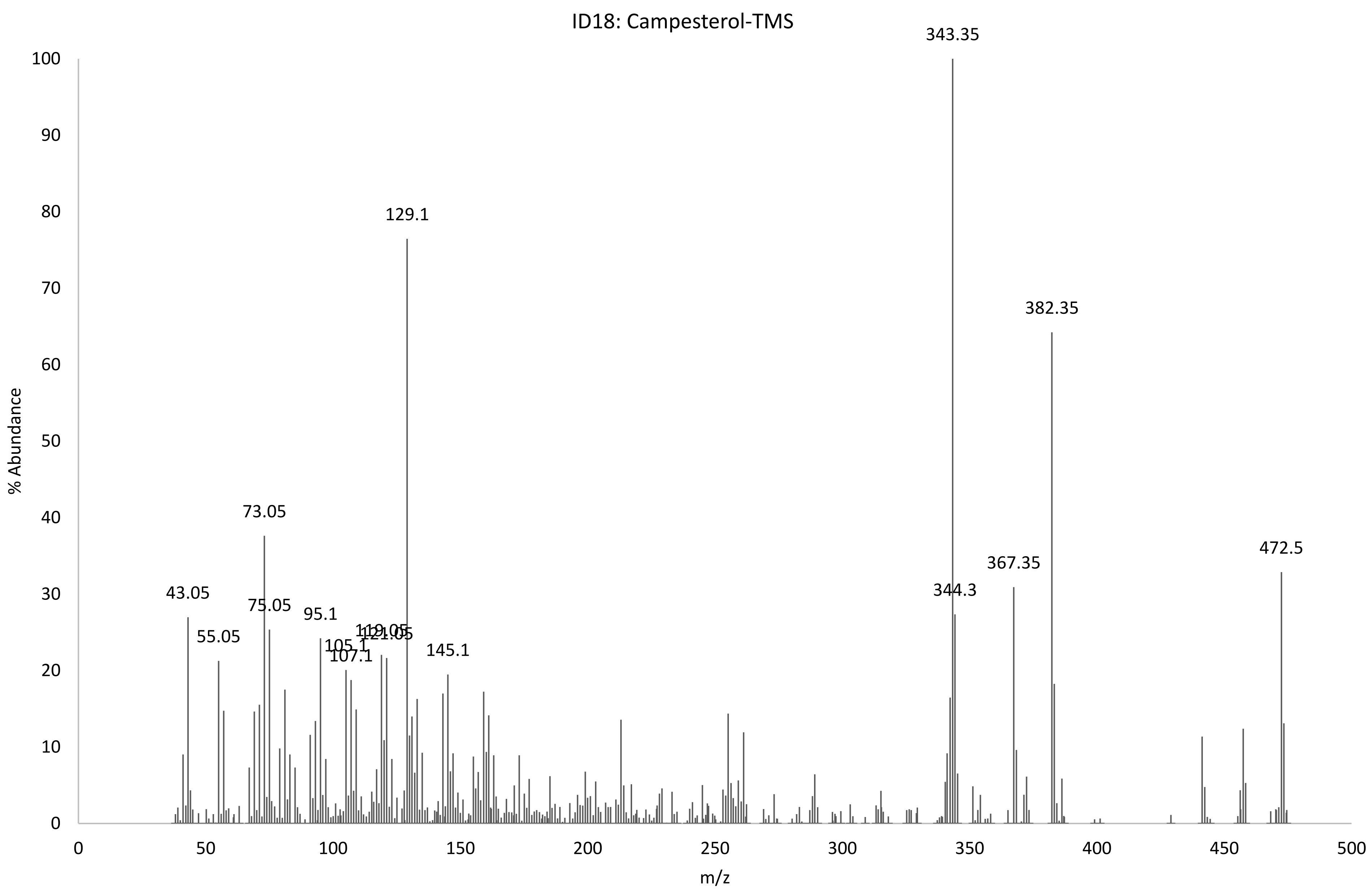
ID18: Campesterol-TMS
% Abundance
m/z
343.35
129.1
382.35
73.05
472.5
367.35
43.05
344.3
75.05
95.1
55.05
119.05
121.05
105.1
107.1
145.1

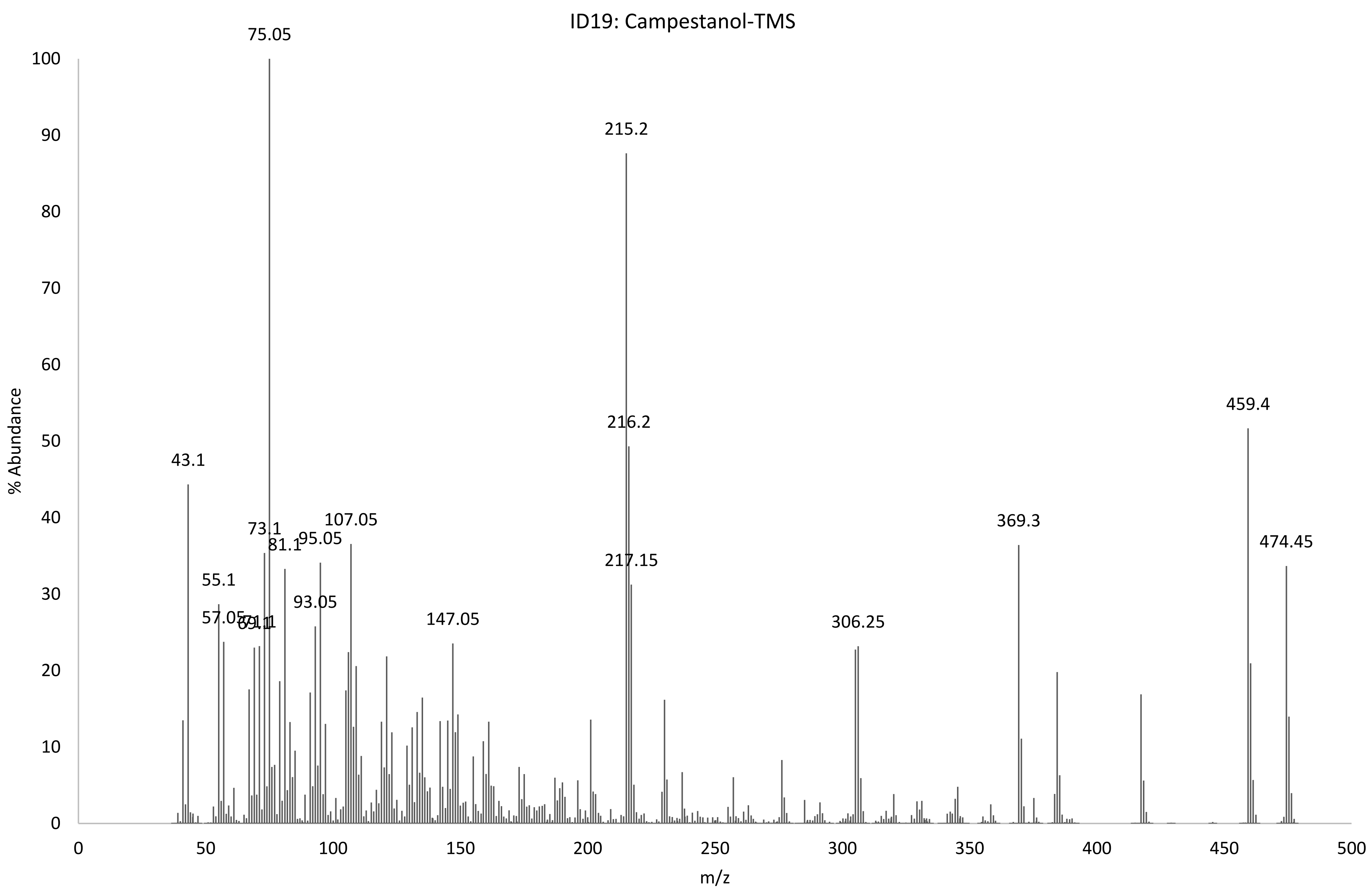
ID19: Campestanol-TMS
% Abundance
m/z
75.05
215.2
216.2
217.15
459.4
474.45
43.1
73.1
81.1
95.05
107.05
55.1
57.05
93.05
147.05
306.25
369.3
0
10
20
30
40
50
60
70
80
90
100
0
50
100
150
200
250
300
350
400
450
500

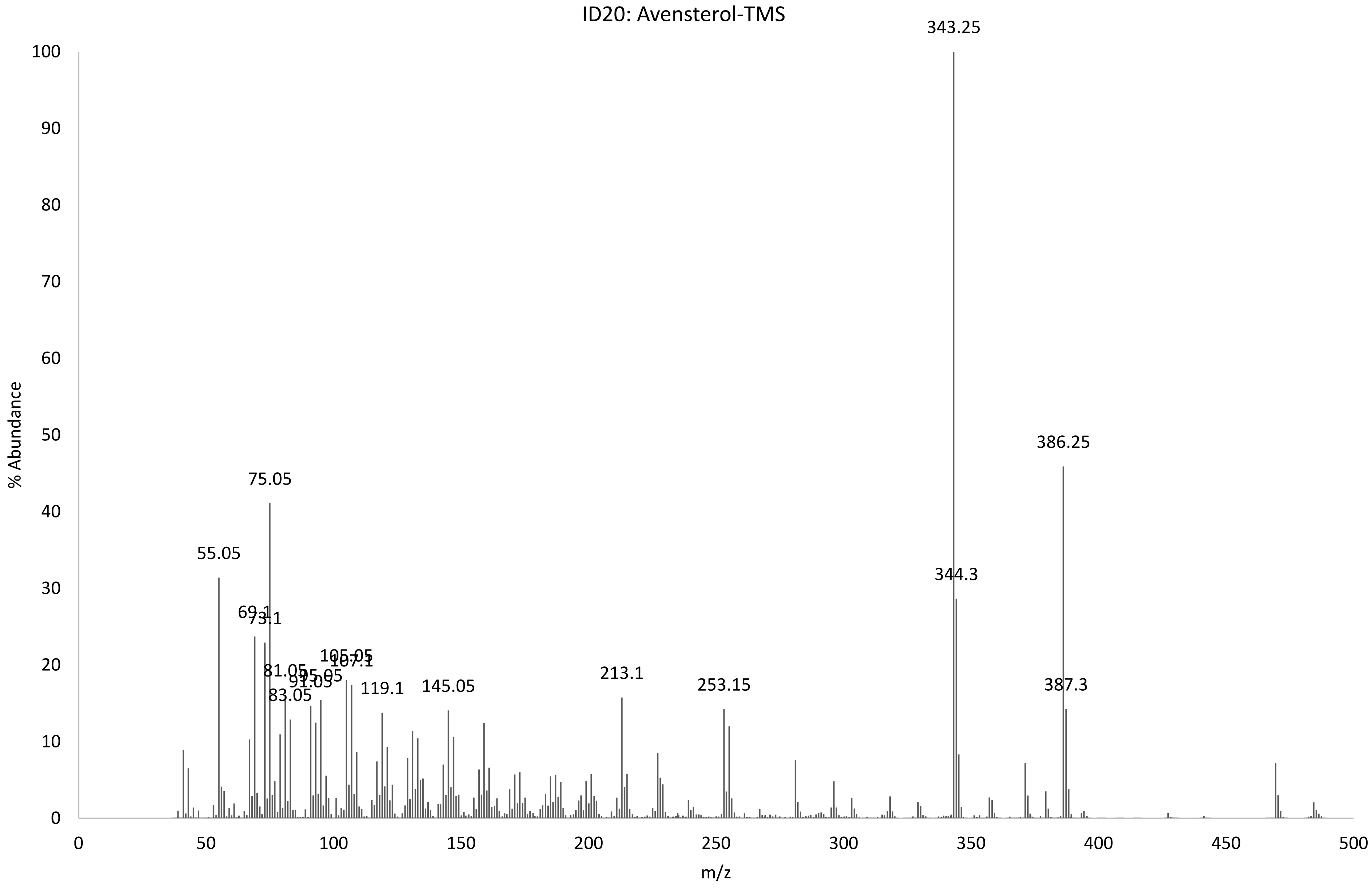
ID20: Avensterol-TMS
% Abundance
m/z
0
10
20
30
40
50
60
70
80
90
100
0
50
100
150
200
250
300
350
400
450
500
343.25
386.25
75.05
55.05
344.3
69.1
73.1
105.05
107.1
81.05
95.05
91.05
83.05
119.1
145.05
213.1
253.15
387.3

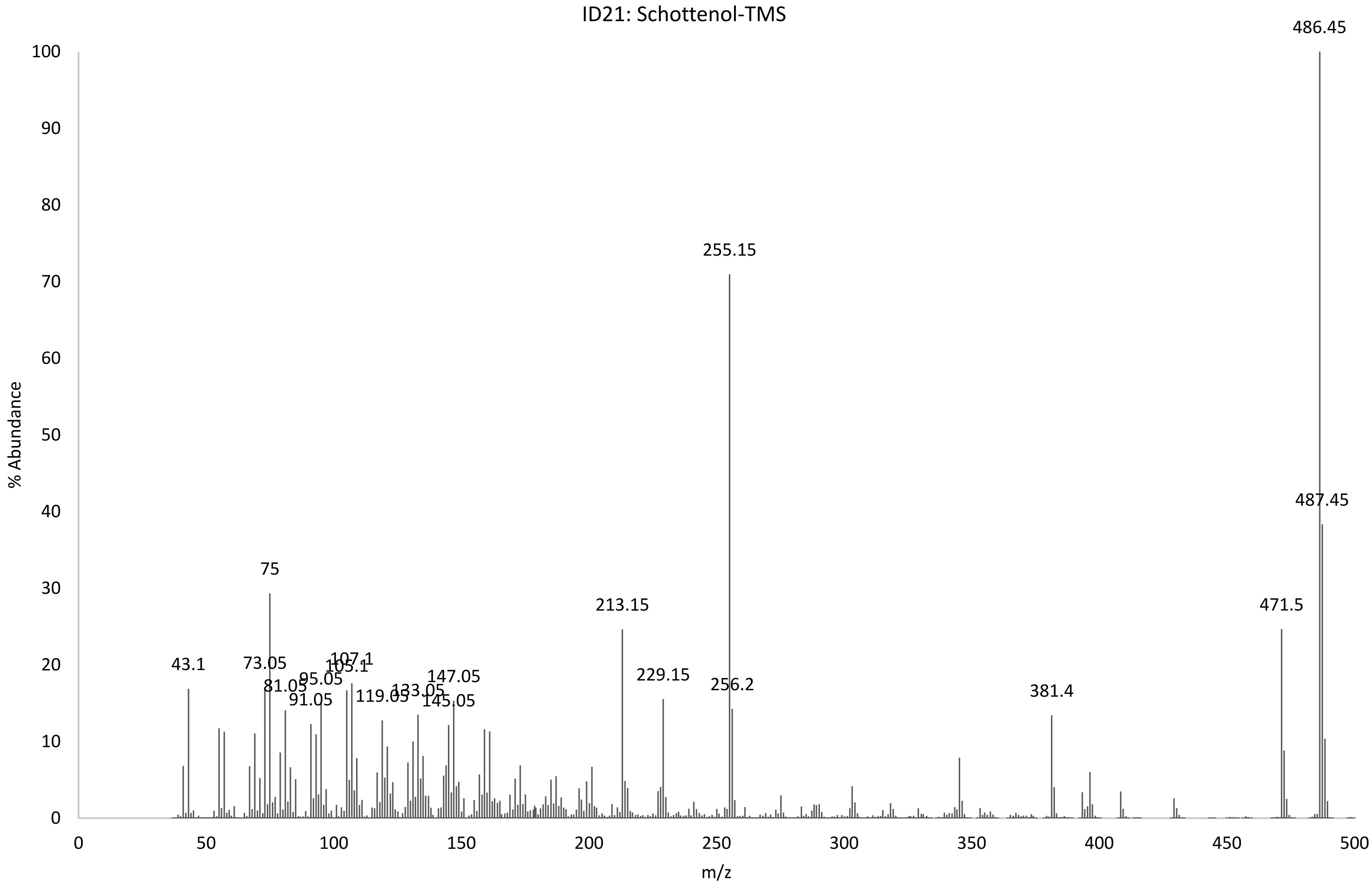
ID21: Schottenol-TMS
% Abundance
m/z
0
10
20
30
40
50
60
70
80
90
100
0
50
100
150
200
250
300
350
400
450
500
43.1
73.05
75
81.05
91.05
95.05
105.1
107.1
119.05
133.05
145.05
147.05
213.15
229.15
255.15
256.2
381.4
471.5
486.45
487.45

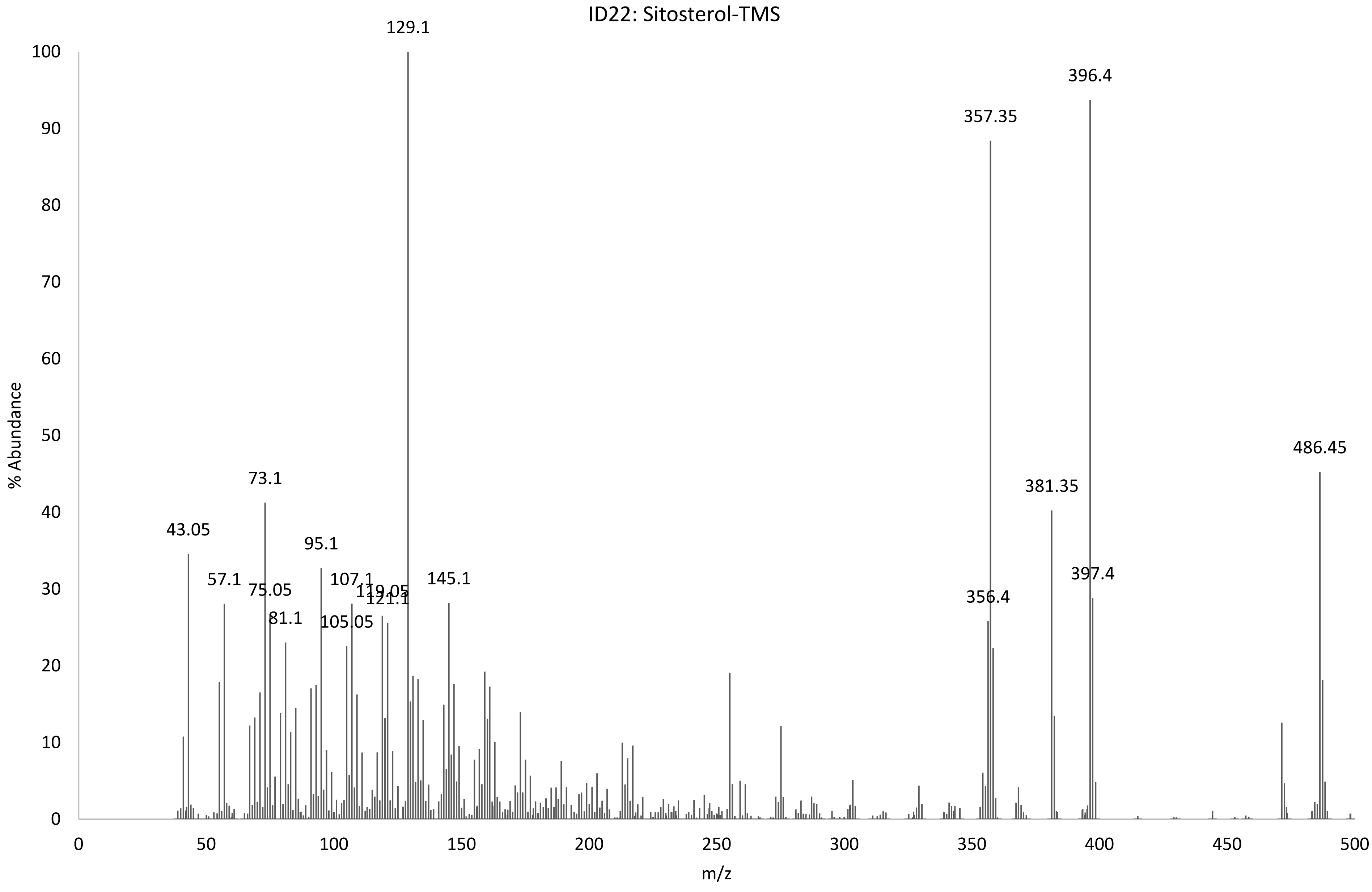
ID22: Sitosterol-TMS
% Abundance
m/z
129.1
396.4
357.35
486.45
381.35
43.05
73.1
95.1
57.1
75.05
81.1
105.05
107.1
119.05
121.1
145.1
356.4
397.4

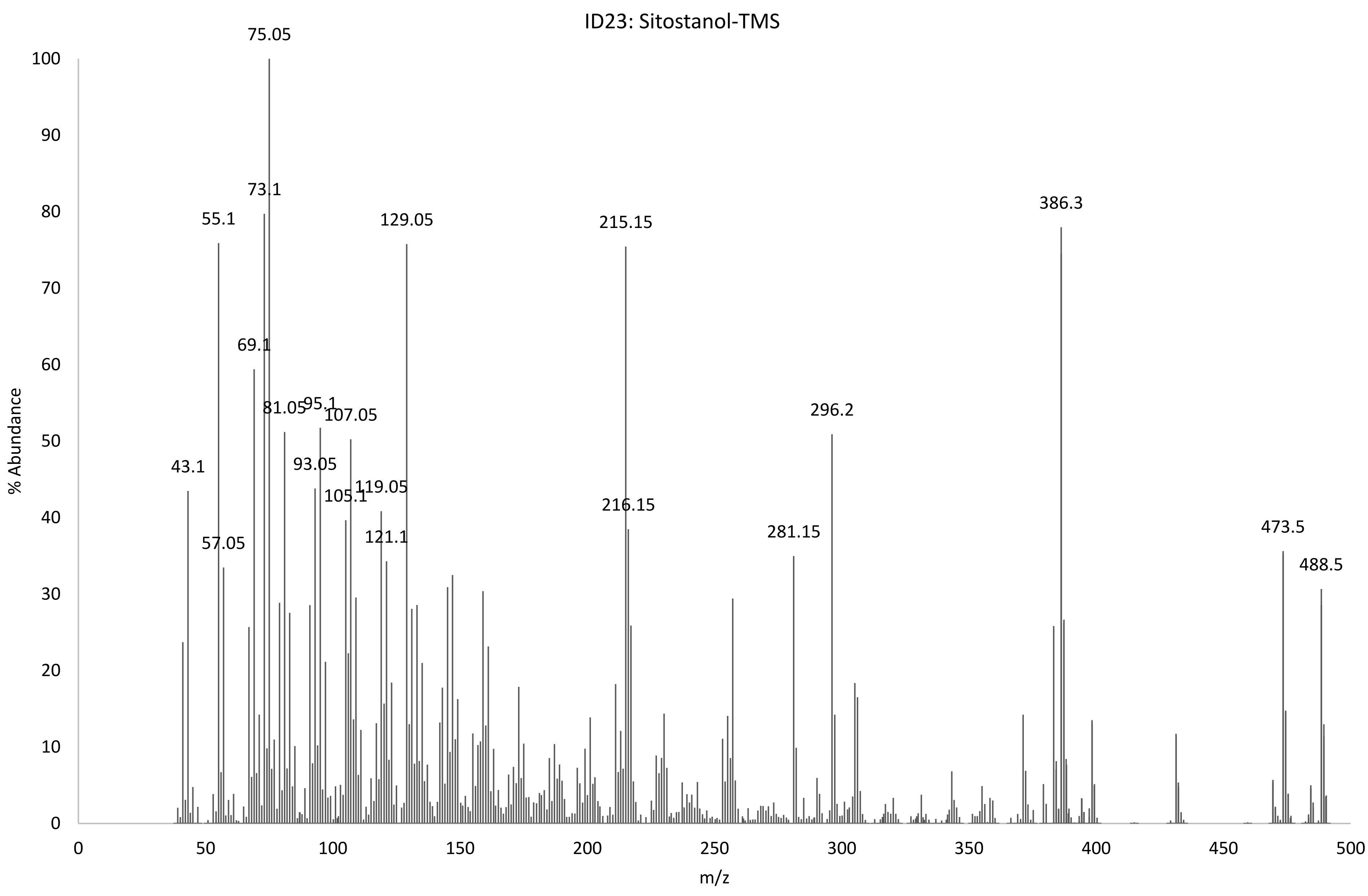
ID23: Sitostanol-TMS
% Abundance
m/z
75.05
73.1
55.1
129.05
215.15
386.3
69.1
81.05
95.1
107.05
93.05
105.1
119.05
121.1
43.1
57.05
216.15
281.15
296.2
473.5
488.5

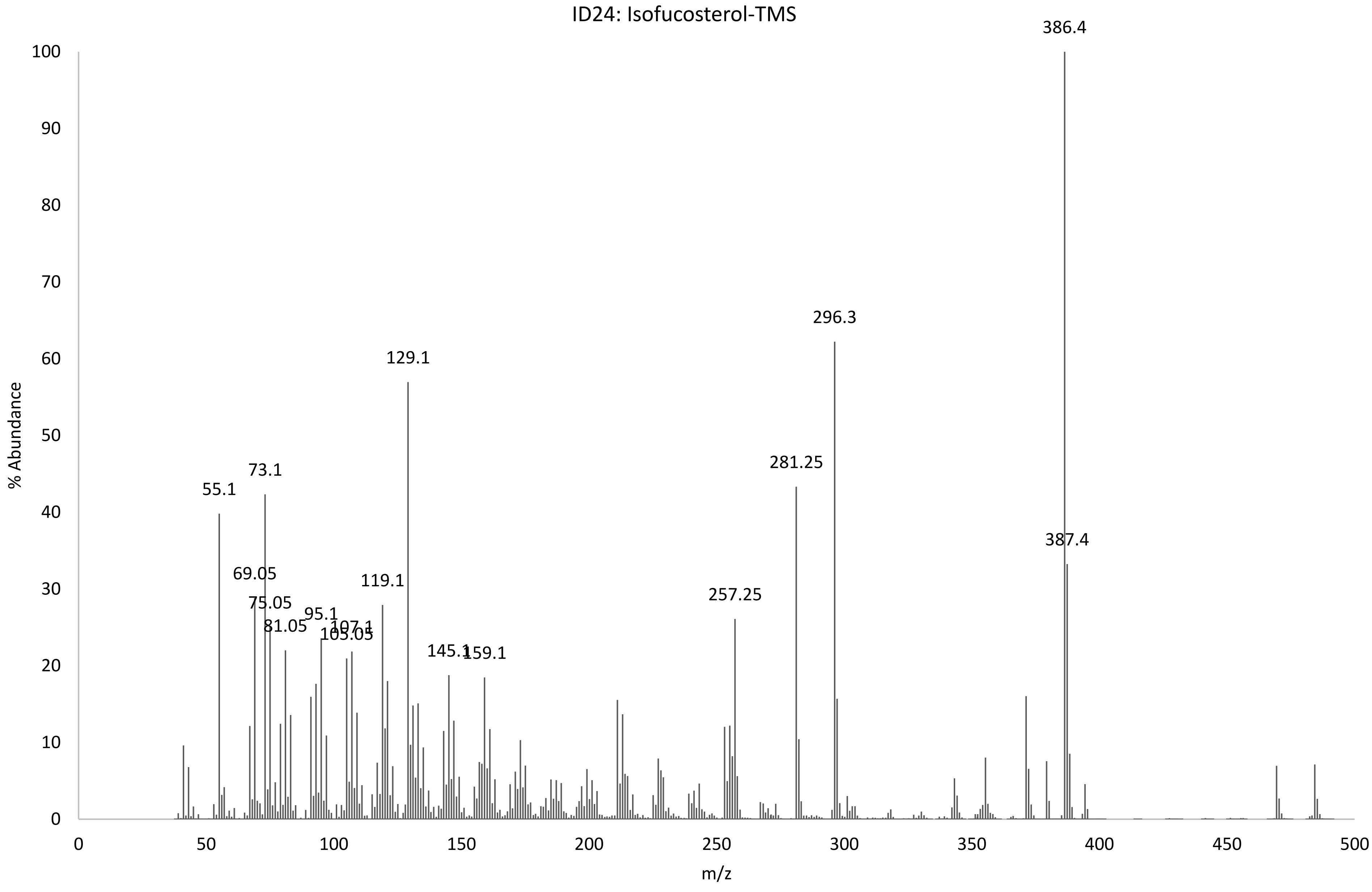

ID24: Isofucosterol-TMS
% Abundance
m/z
386.4
296.3
129.1
281.25
73.1
55.1
387.4
69.05
119.1
257.25
75.05
95.1
81.05
107.1
105.05
145.1
159.1
0
10
20
30
40
50
60
70
80
90
100
0
50
100
150
200
250
300
350
400
450
500

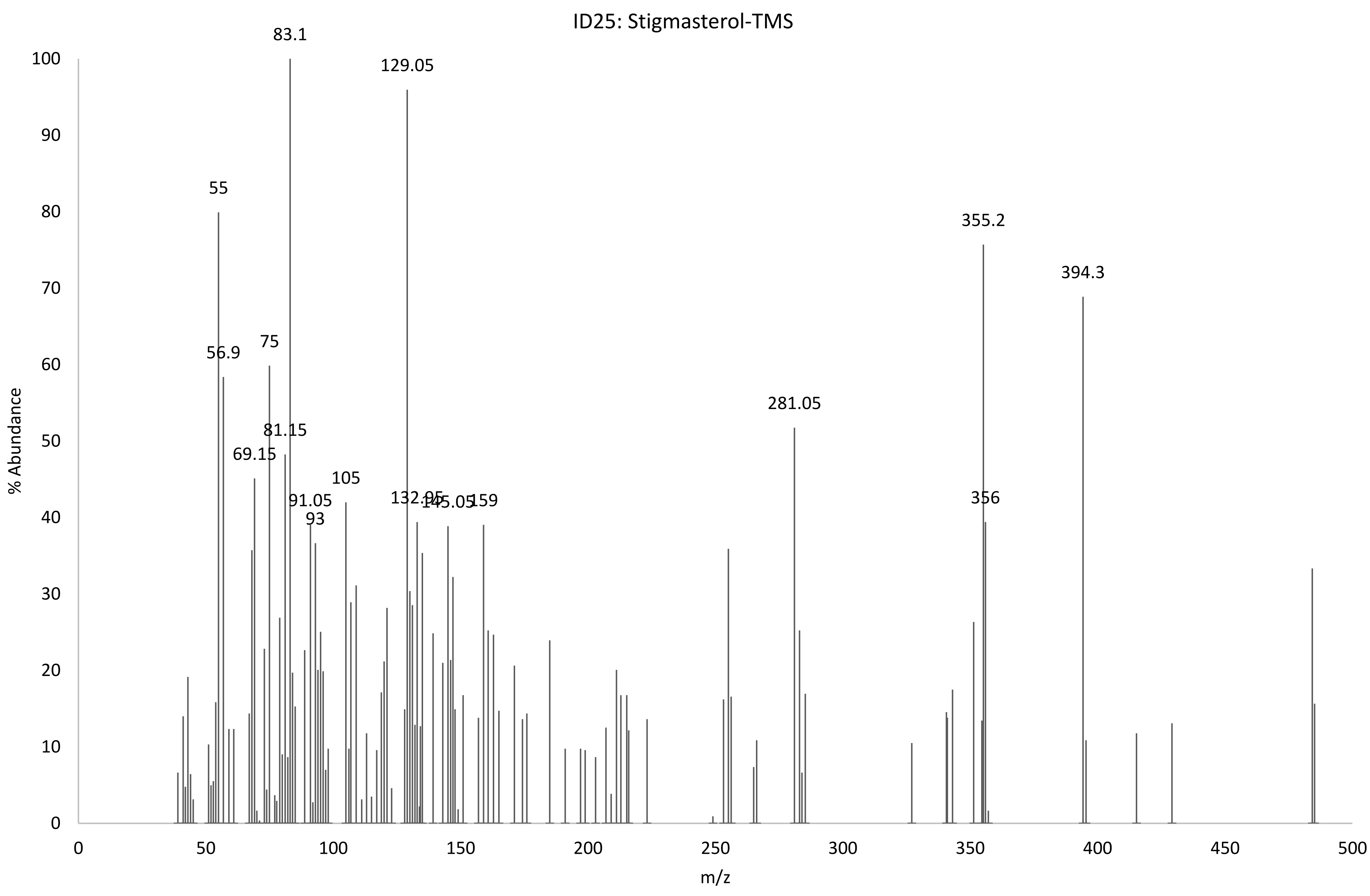
ID25: Stigmasterol-TMS
% Abundance
m/z
0
10
20
30
40
50
60
70
80
90
100
0
50
100
150
200
250
300
350
400
450
500
83.1
129.05
55
56.9
75
81.15
69.15
105
91.05
93
132.05
145.05
159
281.05
355.2
356
394.3

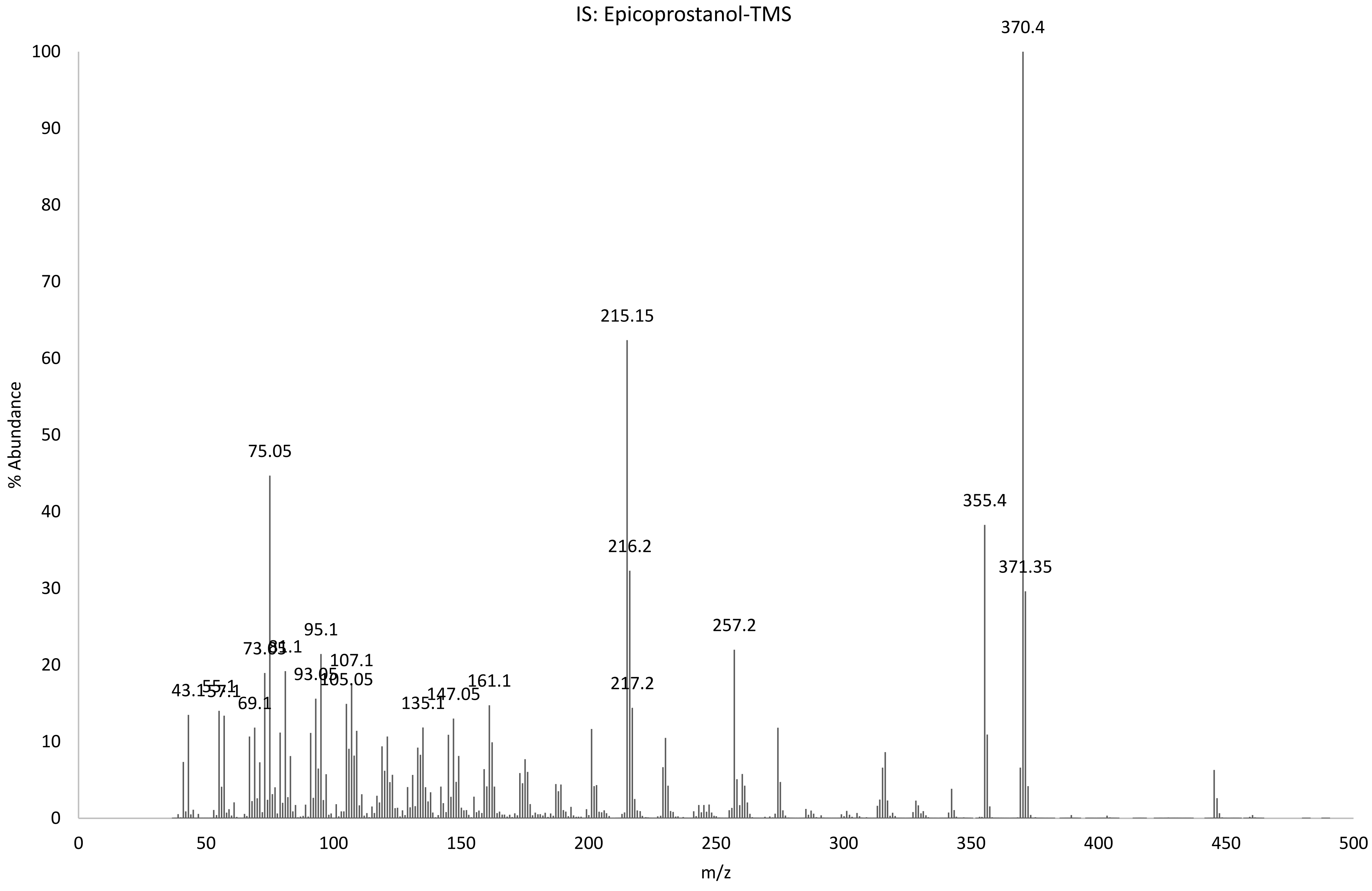
IS: Epicoprostanol-TMS
370.4
215.15
75.05
355.4
216.2
371.35
257.2
73.05
81.1
95.1
107.1
93.05
105.05
43.1
55.1
57.1
69.1
135.1
147.05
161.1
217.2
% Abundance
m/z
0
10
20
30
40
50
60
70
80
90
100
0
50
100
150
200
250
300
350
400
450
500
